# Aberrant development of glutamatergic inputs to PV and SOM interneurons shape amygdala excitability and oscillatory dynamics after early life stress

**DOI:** 10.1101/2025.08.30.673226

**Authors:** Vasilii Shteinikov, Jun Kyu Rhee, Simo Ojanen, Ada-Julia Kunnari, Joni Haikonen, Maxime Picard, Otto Litkey, Anton Lakstygal, Henrike Hartung, Sari E. Lauri

## Abstract

Early-life stress (ELS) induces persistent amygdala dysfunction and affects amygdala-related emotional behaviors, yet the developmental and physiological mechanisms driving these effects remain poorly understood. Here, we provide a comprehensive electrophysiological characterization of the effects of ELS on parvalbumin and somatostatin interneurons (INs) as well as principal neurons (PNs) in the mouse lateral amygdala (LA) across development. Additionally, we correlate these findings to activity of the LA circuitry *in vivo*, using Neuropixels recordings in awake mice. In preweaning juveniles, the effects of ELS were remarkably similar in males and females, involving reduced IN excitability, elevated glutamatergic input to INs and shift in the PN E/I balance. While IN function was largely normalized in adult females, males developed a distinct pathological phenotype characterized by reduced glutamatergic input to INs, impaired recruitment of INs and hyperexcitability of PNs. This male-specific dysfunction correlated with aberrant LA oscillatory dynamics in awake mice and deficits in fear processing. Our data suggest that the impaired glutamatergic wiring of interneurons is a key mechanism underlying the aberrant circuit dynamics in the LA after ELS exposure and contribute to the ELS-induced defects in fear processing. These findings highlight sex-specific developmental trajectories of interneuron connectivity as a potential factor contributing to ELS-induced psychiatric vulnerability.

## INTRODUCTION

Early life stress (ELS) is a recognized risk factor that is associated with heightened vulnerability to psychopathology across the life course. The effects of ELS are particularly pronounced in the amygdala -associated circuits that underlie emotional behaviors and develop during an extended period postnatally. Accumulating data from both humans and animal models highlight amygdala hyperexcitability after ELS, a feature that has been suggested to have a key contribution to pathological anxiety (Rau et al., 2015; Krugers et al., 2016; Guadagno et al., 2021; Tottenham and Sheridan, 2010).

The neurobiological mechanisms underpinning stress-induced amygdala hyperexcitability have been widely studied in adult rodents (reviewed by Zhang et al., 2021). However, as the physiological mechanisms triggering the stress response as well as the responding neural circuits are still immature during the ELS exposure (van Bodegom et al., 2017), the mechanistic data obtained from adults are not directly applicable to ELS. Furthermore, immature networks are highly susceptible to homeostatic plasticity, potentially masking functional changes induced by the early stress by restorative alterations, aimed to balance the overall excitability of the circuit (e.g. Turrigiano & Nelson, 2004; Huupponen et al., 2007). Hence, in order to understand how the early stress exposure leads to adult dysfunction, it is crucial to recognize which cell types and circuit mechanisms are targeted directly by the stress exposure early in life and which are attributable to compensatory plasticity during maturation of the circuitry. This knowledge is also critical for the design of therapeutic strategies to prevent the ELS induced neuropathology before the emergence of the aberrant behaviors.

The existing literature indicates that changes in amygdala reactivity and morphology, including alterations in both principal neurons as well as GABAergic neurons, can be detected in preweaning rodents (P13-22) within two weeks after the ELS exposure (Raineki et al., 2019; Guadagno et al., 2018; Guadagno et al., 2020; Manzano Nieves et al., 2020; Haikonen et al., 2022). The effects of ELS on the GABAergic system are complex, and a mosaic of region-specific, sex-dependent and developmentally transient effects on GABAergic neurons, particularly on the parvalbumin - expressing subtype of GABAergic interneurons (PV INs), have been reported (Guadagno et al., 2020; Manzano Nieves et al., 2020; Haikonen et al., 2022; Perlman et al., 2021; Woodward and Coutellier, 2021). Most of these studies have focused on morphological and neurochemical maturation of the INs, while their physiological functions within the amygdala microcircuitry in ELS-exposed animals are less well understood. Since PV INs are powerful regulators of the local network activity, their malfunction after ELS exposure could have profound effects on network states implicated in cognitive processing (Akam and Kullman 2014). Accordingly, ELS has immediate effects on functional dynamics of the corticolimbic circuits in rodents (Reincke and Hanganu-Opatz, 2017; Raineki et al., 2019; Haikonen et al., 2022; Donati et al., 2024). Yet, it is not clear how these ELS-induced mechanisms, from individual cells to circuits, are integrated to produce persistent amygdala dysfunction.

In order to understand how the amygdala pathophysiology develops after ELS exposure, we have here studied the effects of ELS on the physiological properties of defined cell types in the mouse lateral amygdala (LA) at two stages of development, pre-weaning juveniles (P18-21) and adults (P60-80) of both sexes. Our data highlight the role of sex-specific, post-insult developmental events in generation of perturbed circuit dynamics that are associated with adult psychopathology after ELS.

## RESULTS

### ELS induced by combined LBN and MS results in impaired fear conditioning and male-specific anxiety – like behavioral features in adulthood

To model ELS, the mouse litters were housed in limited bedding and nesting (LBN) conditions between postnatal days (P)4-14, with an additional unpredictable stress component induced by 1h long maternal separations (MS) at P8, P10 and P12 (from here on, referred to as ELS) (Rice et al., 2008; Johnson et al., 2018) (Figure 1A). To validate the model in our hands, we first monitored the behavior of the dams nursing their litters under standard and ELS conditions at P5 and P10. Consistent with previous reports (Walker et al., 2017), ELS resulted in fragmented care, observed as significantly higher number of nest entries and exists by the dams in ELS conditions compared to controls (Figure 1B). Also, the weight gain of the pups exposed to ELS was significantly lower compared to controls (Figure 1C).

**Figure 1.**
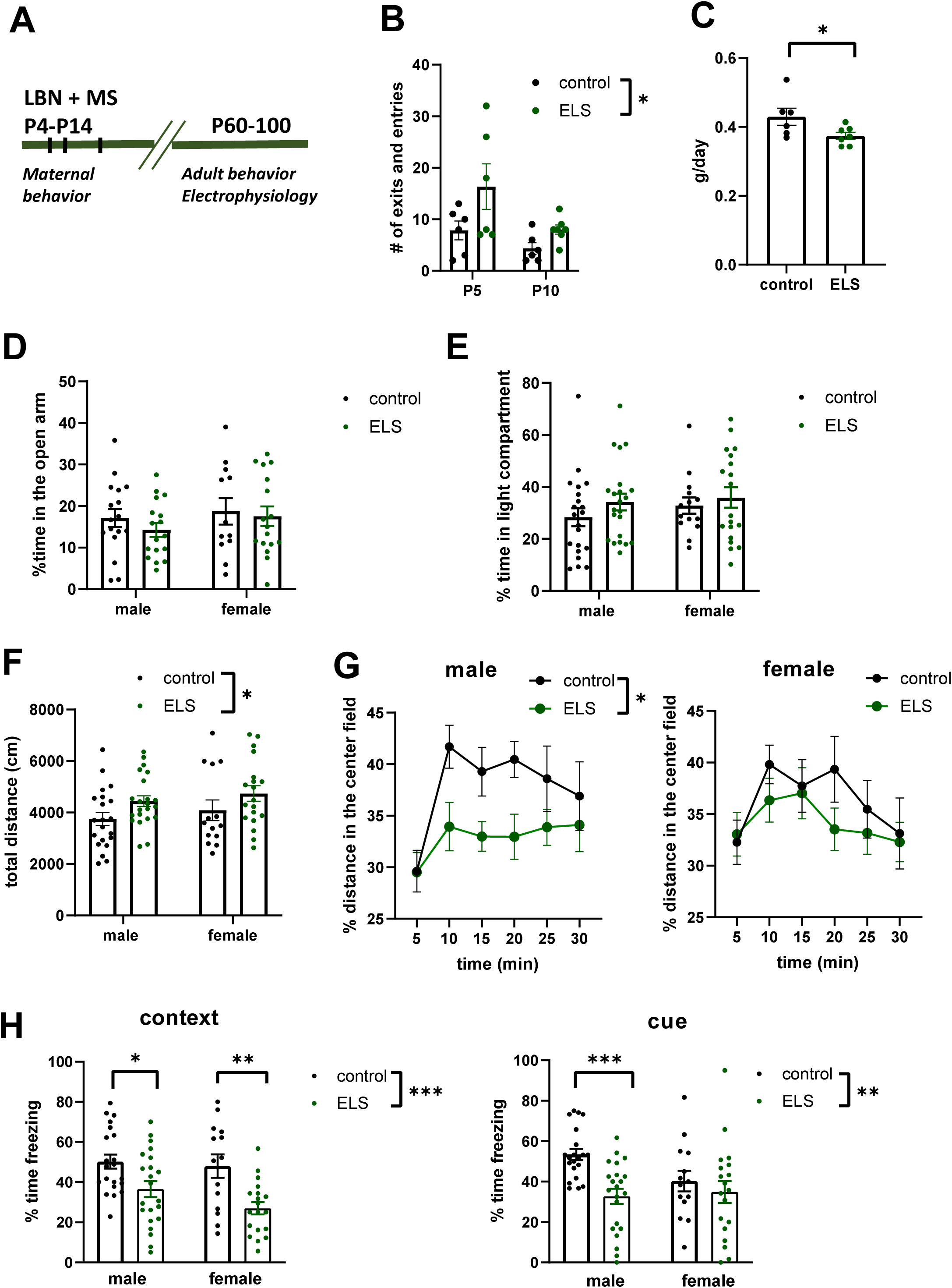
Early life stress (ELS) induced by limited bedding and nesting (LBN) combined with maternal separation (MS) affects fear-related and cognitive behaviors. *A.* Experimental design. Male and female mice were reared under LBN conditions between P4-14 and exposed to MS (1 hour) at P8, P10 and P12. Maternal behavior was followed during the ELS exposure. The behavior of the ELS-exposed pups was analyzed > P60. *B.* Quantified data on dam’s exists and entries to the nest, under control and ELS conditions at P5 and P10. Data is obtained from dams of 6 control and 7 ELS litters. * p=0.03, F _(1, 11)_ = 6.10; RM ANOVA. *C.* The daily weight gain of the pups reared under control and ELS conditions. The pups were weighed at P4 and P14. The individual dots refer to the average weight of all pups (regardless of sex) in each litter (control n=6, ELS n=7 litters). *p=0.02, t=2.70, df=11, unpaired t-test. *D.* Elevated plus maze (EPM) test of the control and ELS exposed mice. The % spent in the open arm of the EPM, for 24 male and 18 female controls, 19 male and 15 female ELS exposed mice. ELS treatment had no effect on the time spent in the open arms (F _(1, 59)_ = 0.76, p=0.39; 2-way ANOVA). *E.* Light-dark compartment test. ELS treatment had no effect on the time spent in the light compartment (F _(1, 72)_ = 1.55, p=0.22, 2-way ANOVA). *F.* OF test. The total distance travelled during the test was higher in the ELS group compared to controls (F_(1, 72)_ = 5.59, *p=0.021, 2-way ANOVA). *G.* The time spent in the center field of the OF arena, for control and ELS exposed males and females. Males, F _(1, 41)_ = 5.27, *p=0.027; females F _(1, 31)_ = 0.63, p=0.43, RM ANOVA). *H.* Classical fear conditioning. Contextual fear conditioning was impaired in both male and female, ELS exposed mice (F _(1, 72)_ = 18.30, *** p<0.0001, 2-way ANOVA; Tukey’s test for males, *p = 0.012; females **p=0.002). ELS treatment also affected cue-dependent fear conditioning (F _(1, 72)_ = 9.354, **p=0.0031, 2-way ANOVA), predominantly in males (Tukey’s test for males, ***p = 0.0008; females p=0.403). All data represent mean ± s.e.m.

Adult mice that had experienced ELS were exposed to standard tests probing hippocampus and amygdala dependent behaviors. We observed no differences between the ELS and control groups in the elevated plus maze or in light-dark compartment tests (Figure 1D,E). However, in the open field, the ELS-exposed mice were slightly more active compared to controls, based on the total distance travelled during the test (Figure 1F). In addition, the males, but not females, avoided the central zone of the open field arena (Figure 1G). Furthermore, the performance of the ELS-exposed mice was significantly impaired in the classical fear conditioning test. While both sexes displayed impaired contextual fear memory (Figure 1H), only ELS males showed impaired cue-dependent fear memory (Figure 1 H).

These data indicate that exposure to ELS, composed of combined LBN and MS, results in altered fear-related cognitive behaviors, particularly in male mice. While the contextual fear conditioning was strongly impaired after ELS in both sexes, only ELS males showed impaired cued fear conditioning and anxiety-like behavior in the open field test.

### ELS induces sex-dependent changes in principal neuron excitability and synaptic E/I balance in the LA

After validating that ELS resulted in alterations in amygdala dependent behaviors, we went on to investigate associated functional changes in the LA microcircuits. First, we performed patch clamp recordings from LA principal neurons (PNs) in acute slices from male and female control and ELS-exposed mice and investigated the effects of ELS on their functional properties across development.

The excitability of the LA PNs was assessed by testing the action potential (AP) firing frequency in response to depolarizing current steps. In juveniles, AP frequency was significantly higher in ELS females compared to controls but not affected by ELS in males (Figure 2A). In the adults, AP frequency was higher in ELS exposed mice compared to controls (F_(1, 42)_ = 5.88, p=0.020), consistent with amygdala hyperexcitability. This effect was prevalent in males but did not reach statistical significance in females (Figure 2B). Together, these data show that the LA PN hyperexcitability after ELS exposure develops in a sex-dependent manner, being observed in juvenile females, but emerging only later in life in males.

**Figure 2.**
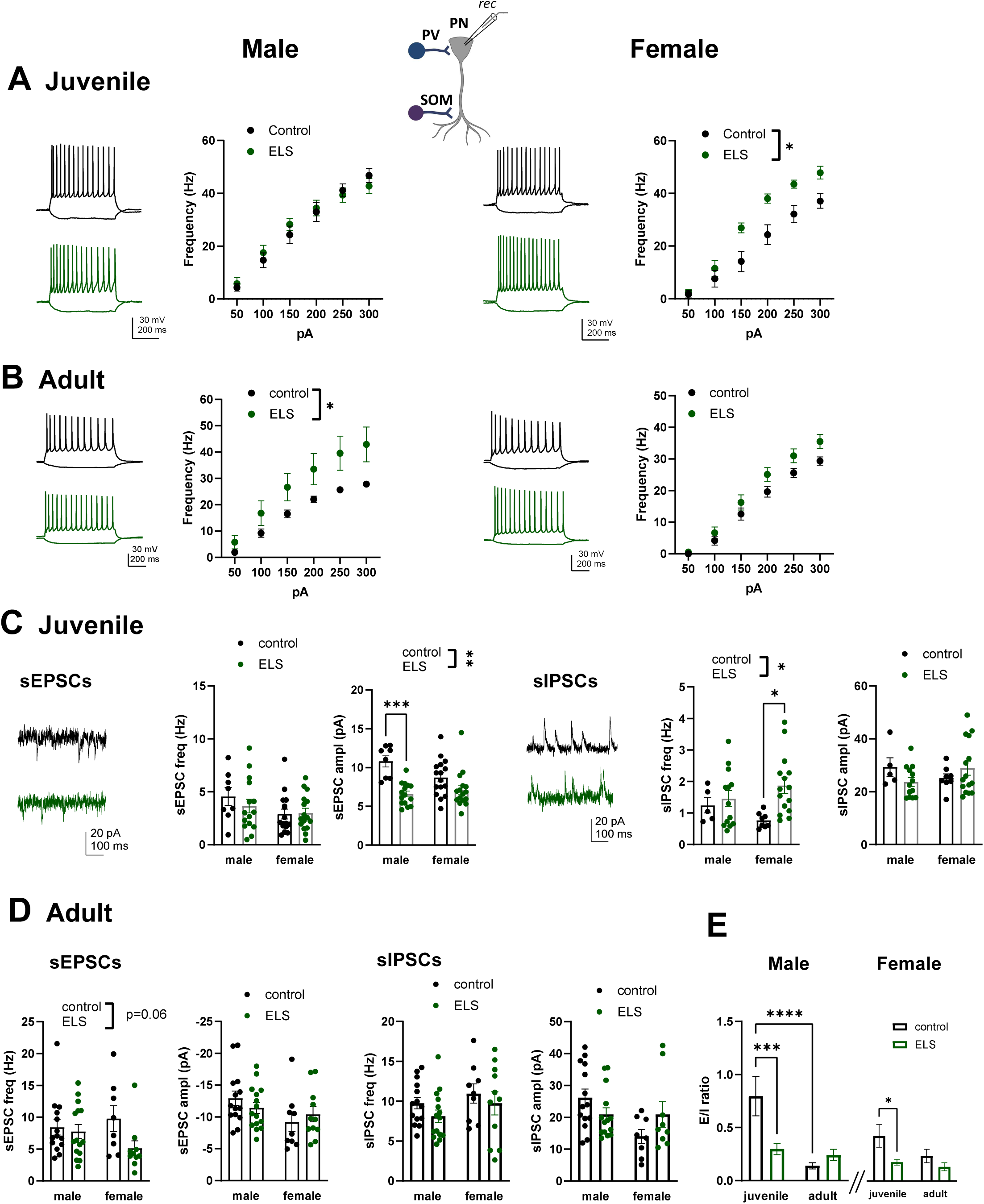
ELS influences excitability and synaptic input of LA principal neurons in a developmentally regulated and sex -specific manner A. LA PN firing rate in response to depolarizing current steps in control and ELS-exposed juvenile mice. Example traces in response to 150 pA steps are shown in the left. Pooled data is collected from males [control n = 14(3), ELS n = 11(3)] and females [control n = 10(3), ELS n = 12(3)]. The n numbers here and all the following figures refer to number of recorded cells, with the number of animals in parentheses. * p=0.032, F_(1, 11)_ = 6.071, RM ANOVA. B. Similar data as in A, for adult mice. Male control n=12(3), ELS n=14(3); Female control n= 15(3), ELS n=15(3). *p=0.043, F_(1, 21)_ = 4.33, RM ANOVA. C. Synaptic inputs to LA PNs, in control and ELS-exposed juvenile mice. Example traces show recordings of sEPSCs and sIPSC from male control (black) and ELS (green) slices. Pooled data for sEPSCs: male control n=8(2), ELS n=15(3); female control n=16(3), ELS n=16 (3); **p < 0.0001, F_(1,51)_ = 23.75, 2-way ANOVA; ***p=0.0002, Tukey’s test. sIPSCs: male control n=5(2), ELS n=13(3); female control n=8(3), ELS n=15(3). *p=0.022, F_(1,37)_ = 5.678, 2-way ANOVA; *p=0.018 for Tukey’s post-hoc test in females) D. Similar data as in C, for adults. sEPSCs: male control n=14 (8), ELS n=15 (6); female control n=9 (6), ELS n=10 (6); sIPSCs: male control n=14 (8), ELS n=15(6); female control n=9(6), ELS n=11(6). E. Development of synaptic E/I ratio in control and ELS-exposed, male and female mice. The synaptic E/I ratio is calculated as the charge ratio of the spontaneous glutamatergic / GABAergic synaptic events to individual cells from the data shown in panels C and D. Synaptic E/I ratio was significantly affected by age and ELS in males (age, F_(1, 37)_ = 24.73, p<0.0001; ELS, F_(1, 37)_ = 7.64, p=0.009, 2-way ANOVA) and by ELS in females (age, F_(1, 37)_ = 4.06, p=0.051; ELS, F_(1, 37)_ = 9.25, p=0.0043). Effect of ELS in juvenile males, *** p=0.0006; females *p=0.021; across development in control males, ****p<0.0001, Holm-Sidak.

Analysis of the passive membrane properties revealed no differences between the control and ELS groups in juveniles, however, in the adults ELS exposure was associated with small, sex-independent elevation in the resting membrane potential (control -64.9 ± 1 mV, ELS -61.6 ± 1 mV; F_(1, 43)_ = 5.63; p=0.022; Supplementary Figure 1A).

We next analyzed synaptic inputs to principal neurons under voltage clamp. In these experiments, the cells were first held at -70 mV to record spontaneous glutamatergic synaptic events (sEPSCs), after which the membrane potential was raised to 0 mV, the reversal potential of glutamate receptor mediated currents, to record spontaneous GABAergic activity (sIPSCs) in the same cell (Figure 2C). ELS treatment had no effect on sEPSC frequency in the juveniles, but sEPSC amplitude was significantly lower in the ELS group compared to controls, particularly in males (Figure 2C). sIPSC frequency was higher in the ELS-exposed juveniles than in controls, particularly in females (Figure 2C). In the adults, no significant differences in the frequency or amplitude of spontaneous synaptic events were detected (Figure 2D).

To estimate the overall impact of the synaptic inputs to the target cell we calculated the synaptic E/I ratio in ELS and control mice across development. This analysis revealed that in control mice, the E/I ratio decreased substantially during development, particularly in males (Figure 2E). Interestingly, in the ELS group, the E/I ratio in juveniles was similar to adults and significantly lower compared to controls in both sexes (Figure 2E). These data suggest that the synaptic E/I balance develops prematurely in the ELS-exposed mice, reaching adult levels already in juveniles.

### Impact of ELS on PV INs: developmental and sex-specific effects on firing properties and glutamatergic synaptic inputs

Previous studies have linked ELS exposure to changes in density of PV INs and PV staining intensity in the amygdala (Guadagno et al., 2020; Manzano Nieves et al., 2020; Haikonen et al., 2022), yet how these findings correlate with physiological functions of PV INs remains unclear. To study how ELS affects physiological properties of PV INs, we made patch clamp recordings from PV INs in acute slices from control and ELS-exposed, PVCre-TdTomato reporter mice. ELS has highly reproducible effects across mouse lines (Molet et al., 2016; Walker et al., 2017). Accordingly, both the PVCre-TdTomato pups reared under ELS conditions gained less weight compared to respective controls (controls, 0.39 ± 0.011 g/day; ELS 0.26 ± 0.012 g/day; p<0.0001, two-tailed t-test).

The AP firing frequency of PV INs in response to depolarizing current steps was significantly lower in ELS exposed juveniles compared to controls (Figure 3 A). These differences were observed in both sexes could be largely attributed to low input resistance and high rheobase in the ELS group (Supplementary Figure 2), features observed in amygdala PV INs also after chronic stress in adults (Ryazantseva et al., 2024). In contrast, no significant differences in the PV IN firing frequency were detected between control and ELS-exposed adults (Figure 3B).

**Figure 3.**
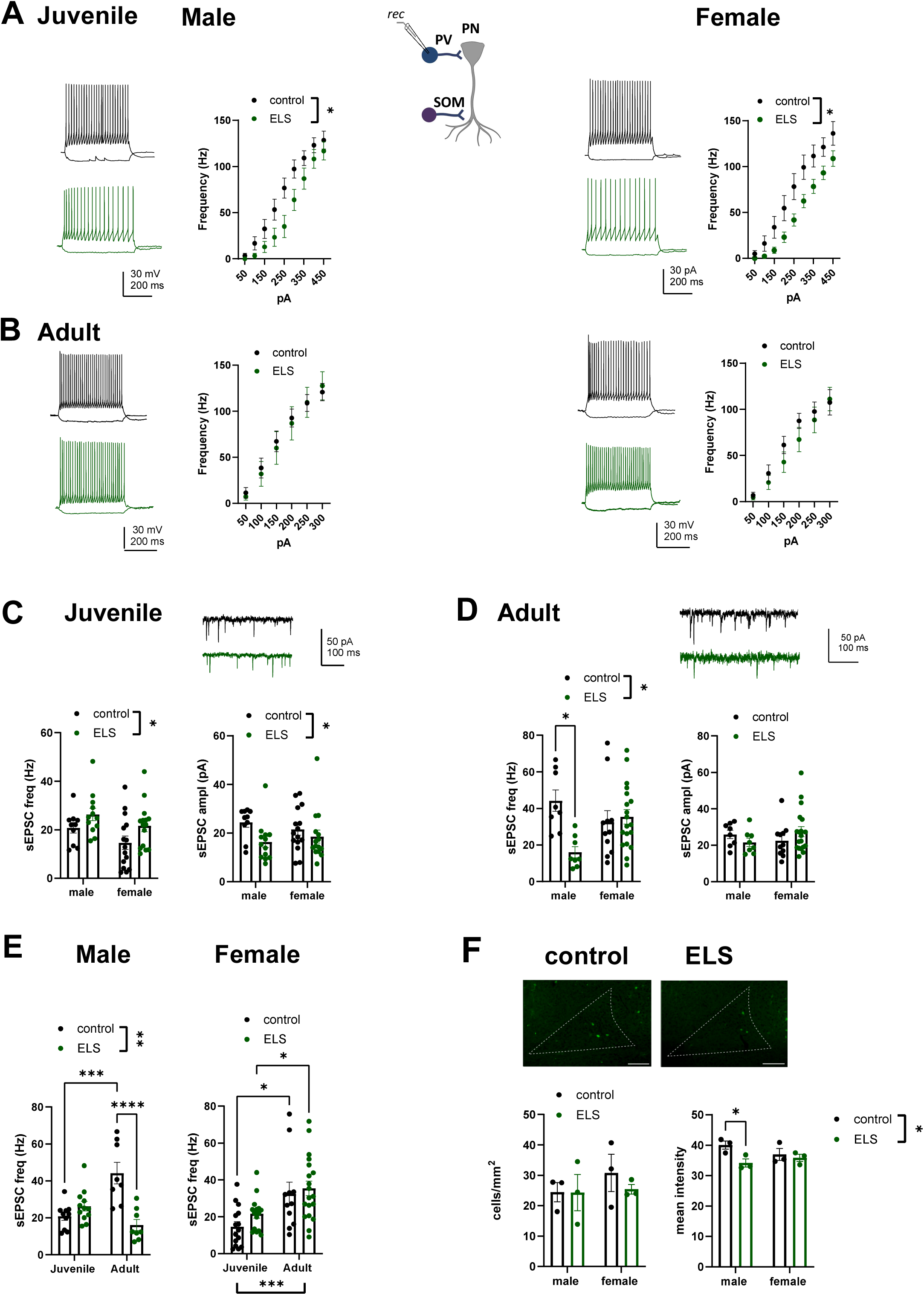
ELS treatment has developmentally transient effects on the firing properties and glutamatergic connectivity of PV INs. A. PV IN firing rate in response to depolarizing current steps in control and ELS exposed juvenile PV-TdTom mice. Example traces in response to 150 pA current steps are shown in the left. Pooled data is collected from males [control 11(3), ELS 14(3), *p=0.048, F_(1, 23)_ = 4.375, RM ANOVA] and females [control n=11(3), ELS 13(3), *p=0.013, F _(1, 22)_ = 7.31, RM ANOVA]. B. Similar data on PV IN firing rates as in A, for adults. Males, control n=19 (6), ELS n=11(5), p=0.80, RM-ANOVA; females, control n=12(4), ELS n=18(7), p=0.44, RM ANOVA. C. Glutamatergic synaptic inputs (sEPSCs) to PV INs in control and ELS exposed juvenile mice. Example traces for sEPSC recordings from male control (black) and ELS (green) slices. Pooled data on the effect of ELS on sEPSC frequency and amplitude for males [control n=10(3), ELS n=12 (3] and females [control n=15(3), ELS n=15 (3)]. The effect of ELS on sEPSC frequency, * p=0.028, F_(1, 49)_ = 5.11; amplitude, * p=0.025, F_(1, 51)_ = 5.30, 2-way ANOVA. D. Similar data as in C, for adults. Example traces from males and pooled data for males [control n=8(2), ELS n=12(3),] and females [control n=11(3), ELS n=15(3)]. sEPSC frequency, * p=0.023, F_(1, 42)_ = 5.548, 2-way ANOVA for all data, *p=0.010, Tukey’s test for males. E. Replotting of the data from C and D, to illustrate development of the sEPSC frequency and amplitude across development in control and ELS exposed males and females. Effect of ELS across development in males F_(1, 34)_ = 10.85, **p=0.0023; ***p=0.0002, **** p<0.0001, Tukey’s test. In females, 2-way ANOVA indicated a significant effect of age (F _(1, 56)_ = 16.09, p=0.0002) but not ELS treatment F _(1, 56)_ = 1.601, p=0.2110; Tukey’s test for controls *p=0.0206, ELS *p=0.0477. F. Parvalbumin immunostaining in the LA of control and ELS exposed juvenile mice. Example images of the staining in males, as well as quantified data on the density of PV labelled neurons and on PV labeling intensity. Data is pooled from at least 8 sections / mouse, and from 3 mice in each group. Effect of ELS in PV intensity *p=0.032, F _(1, 8)_ = 6.73, 2-way ANOVA; * p=0.021, Holm-Sidak. Scale bar is 100 µm.

We also studied the effect of ELS on glutamatergic inputs to PV INs across development. In juvenile PV INs, sEPSC frequency was higher and sEPSC amplitude lower in ELS-exposed mice compared to controls, in both sexes (Figure 3C). In adult PV INs, sEPSC frequency in males, but not females, was lower in the ELS group than in controls (Figure 3D), while no differences between the groups were detected in sEPSC amplitude. In controls, sEPSC frequency increased during development in both sexes, but similar increase was not observed in the ELS males (Figure 3E).

To correlate the functional data to PV expression levels, we analyzed the PV staining intensity in the LA of control and ELS-exposed juvenile mice. The density of PV positive neurons in the LA was not affected by the ELS treatment, however, the mean intensity of PV staining was slightly lower in the ELS group than in controls. Post-hoc tests indicated a significant effect of ELS in males but not in females (Figure 3F).

Together, these data show that ELS affects PV IN firing properties as well as excitatory inputs to PV INs in juveniles. Most of these effects are developmentally transient and normalized towards adulthood. In adults, the predominant difference between the groups was a male-specific loss of glutamatergic synaptic inputs to PV INs in ELS-exposed mice.

### The effect of ELS on the physiological properties of SOM INs is similar to that on PV INs

To understand whether the ELS-associated functional changes were specific to PV INs, we also performed *ex vivo* electrophysiological experiments from somatostatin-expressing INs (SOM INs) in the LA of SST-TdTom reporter mice. ELS treatment had a significant effect on weight gain of the SST-TdTom pups (controls, 0.31 ± 0.014 g/day; ELS 0.20 ± 0.021 g/day; p=0.0003, two-tailed t-test).

Depolarization-induced AP firing frequency of the SOM interneurons was lower in the juvenile ELS group compared to controls, in both males and females (Figure 4A). Similar to the PV INs, this effect was associated with lower input resistance (Supplementary Figure 3). In adults, the AP firing frequency was marginally elevated in ELS group as compared to controls when all the data was pooled together (p=0.061, F_(1, 51)_ = 3.67, RM ANOVA), but not when males and females were analyzed separately (Figure 4B).

**Figure 4.**
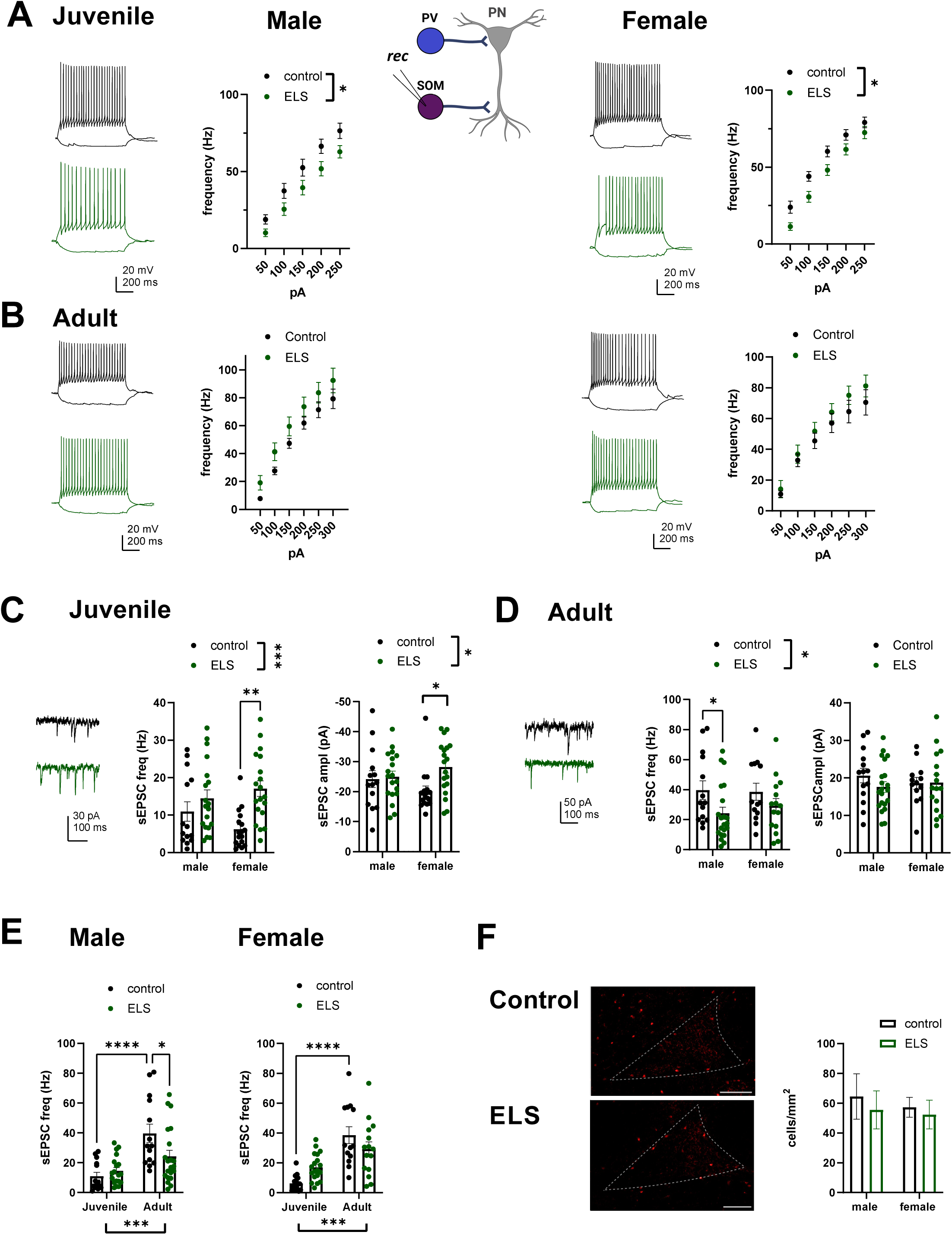
ELS exposure has developmentally transient effects on the firing properties and glutamatergic connectivity of SOM INs. A. SOM IN firing rate in response to depolarizing current steps in control and ELS exposed juvenile SST-TdTom mice. Example traces illustrate the 100 pA depolarizing current step. Pooled data is collected from males [cont n=14(3), ELS n=19 (3), F _(1, 31)_ = 5.00, *p=0.033, RM ANOVA] and females [control n=15 (3), ELS n=18 (3), F _(1, 31)_ = 6.72, *p=0.014, RM ANOVA]. B. Similar data as in A, for excitability of adult SST INs. Males: control n=12(3) , ELS n=14(3); p= 0.11, F_(1, 24)_ = 2.80; females: control n=12(3), ELS n=14(3), p= 0.36, F _(1, 25)_ = 0.88, RM ANOVA. C. Glutamatergic synaptic inputs (sEPSCs) to SOM INs in control and ELS exposed juvenile mice. Example traces for sEPSC recordings from male control (black) and ELS (green) slices. Pooled data on the effect of ELS on sEPSC frequency and amplitude for males [cont n=14(3), ELS n=19(3)] and females [cont n=17(3), ELS n=19 (3)]. The effect of ELS on sEPSC frequency, *** p= 0.0009, F _(1, 65)_ = 12.16; amplitude * p= 0.036, F _(1, 65)_ = 4.59, 2-way ANOVA; post-hoc comparison for sEPSC frequency in females, **p=0.0019, amplitude *p=0.037, Holm-Sidak. D. Similar data as in C, for the effect of ELS on sEPSC in adult SOM INs. Data from males [cont n=14(3), ELS 21(4)] and females [cont n=12(3), ELS n=16(3)]. The effect of ELS on sEPSC frequency, *p= 0.020, F _(1, 59)_ = 5.706, 2-way ANOVA; post-hoc comparison for males, *p=0.030. E. Replotting of the data from C and D, to illustrate development of the sEPSC frequency and amplitude across development in control and ELS exposed males and females. sEPSC frequency was significantly increased during development in both males and females in the control group (males, ***p<0.0001; females, ***p<0.0001; Tukey’s test). In ELS group, sEPSC frequency was not significantly different between juveniles and adults (males, p=0.24; females p=0.065; Tukey’s test). F. Somatostatin immunostaining in the LA of control and ELS exposed juvenile mice. Example images of the staining in males, as well as quantified data on the density of SOM-labelled neurons. Data is pooled from at least 8 sections / mouse, and from 3 mice in each group. No significant differences between groups were detected (p=0.57, F_(1, 8)_ = 0.36, 2-way ANOVA) Scale bar is 100 µm.

Similar to PV INs, ELS treatment affected glutamatergic inputs to SOM INs in a developmentally regulated manner. In the juveniles, both sEPSC frequency and amplitude were higher compared to controls (Figure 4C). In adults, sEPSC frequency was lower compared to controls, and significantly affected in males but not females (Figure 4D). No differences between the groups were detected in sEPSC amplitude. Developmental comparison indicated that sEPSC frequency strongly increased during development of SOM INs in both males and females (p>0.0001, Holm-Sidak)(Figure 4E). This effect was not observed in ELS exposed males and was not significant in ELS females (Figure 4E).

Immunostaining against somatostatin indicated no significant effect of ELS on the density of somatostatin-expressing neurons in the LA (Figure 4F).

Together, these data suggest that ELS has comparable effects the physiological functions of PV and SOM INs in the LA. The excitability of both cell types was lower compared to controls in ELS-exposed juveniles. This effect was sex independent and associated with changes in input resistance. At the same time, the excitatory synaptic drive to PV and SOM INs was elevated in both males and females.

Interestingly, development of glutamatergic inputs to INs seemed to be prematurely arrested in ELS-exposed males, resulting in weak excitatory input to both PV and SOM INs in the adult stage. In adult females, only minor effects of ELS on the PV and SOM INs physiology were observed.

### ELS-exposure influences feedforward inhibition and LTP induction in the adult LA

After characterizing physiological alterations in individual cell types in the LA, we went on to study how these changes affected excitability of the adult LA microcircuit during afferent activation. To mimic activation of cortical inputs, we stimulated the external capsulae and recorded evoked EPSCs (V_hold_ = -70 mV) and disynaptic GABAergic IPSCs (dIPSC, V_hold_ = 0 mV), reflecting activity of the feed-forward inhibitory microcircuits, from the LA principal neurons. The EPSC/dIPSC ratio was higher in ELS treated males compared to controls, while the opposite effect was observed in females (Figure 5A).

**Figure 5.**
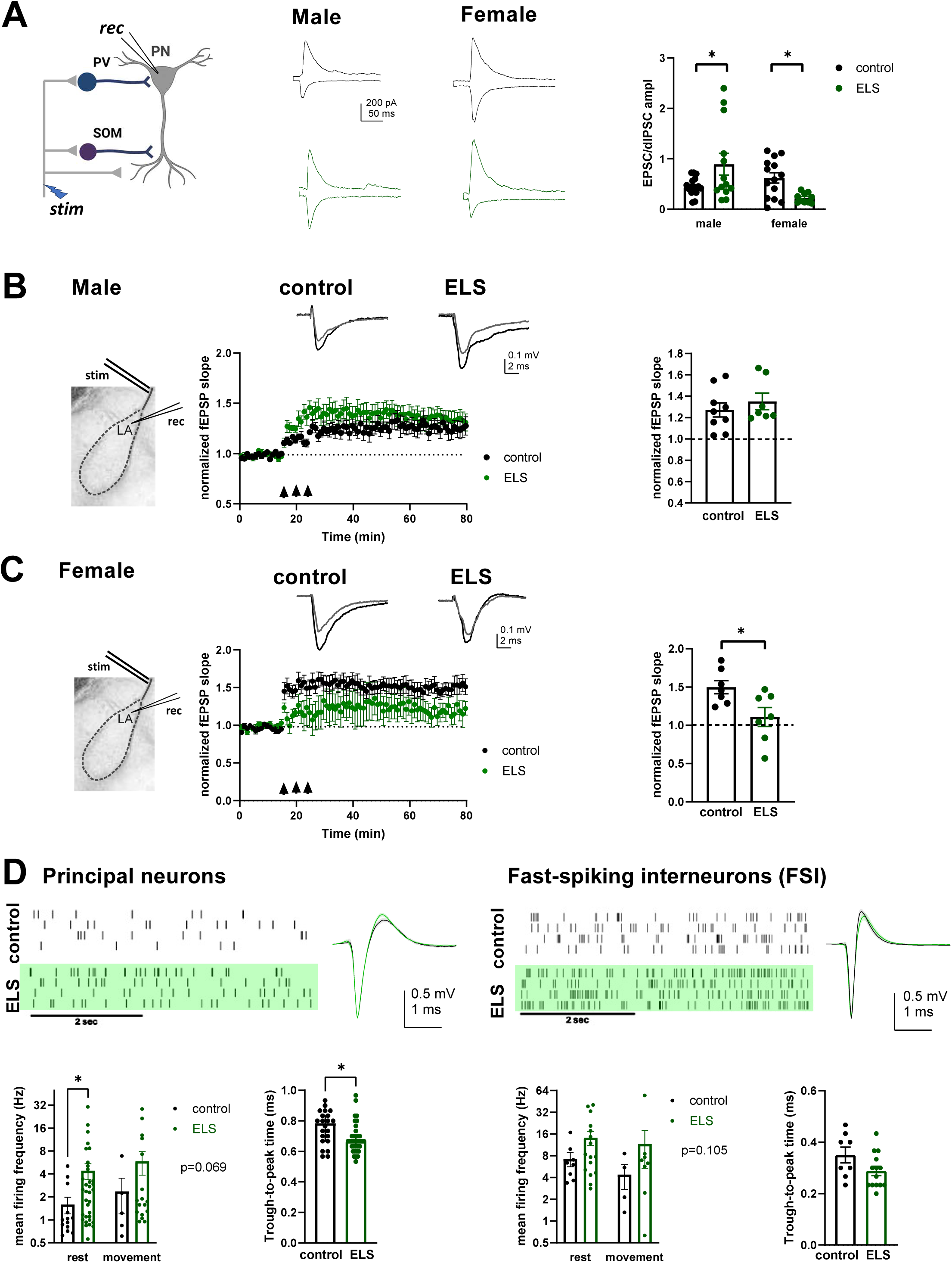
Effect of ELS on excitation-inhibition balance and LTP induction in response to stimulation of cortical afferents in adult LA A. Example traces and pooled data on the balance between excitation (EPSCs) and feedforward inhibition (dIPSCs), recorded from the LA principal neurons in response to stimulation of cortical afferents. EPSC/dIPSC ratio was significantly affected by ELS treatment and sex (interaction, p=0.0011, F_(1, 50)_ = 12.02, 2-way ANOVA) and was higher in ELS males (*p=0.016, Holm-Sidak) and lower in ELS females (*p=0.034, Holm-Sidak) compared to controls. Pooled data from males [control n=17(5), ELS n=13 (3)] and females [control n=13 (4), ELS n=10(3)]. B. A time course plot showing the effect of TBS (arrowheads) on fEPSPs, recorded from LA in response to stimulation of cortical afferents in slices from control and ELS-treated male mice. Superimposed example traces before (black) and 60 min after (grey) TBS are shown on top. Pooled data shows the average level of synaptic potentiation 60 minutes after the TBS in control [n=9(4)] and ELS [n=7(4)] groups. C. Similar data as in B, for the effect of ELS on TBS induced LTP in females. Pooled data for controls, n=7(5) and ELS, n=7(5). * p=0.024, unpaired t-test D. Raster plots and spike waveforms illustrating single-unit firing in the LA of control (black) and ELS-exposed (green) head-fixed awake mice. Pooled data on the mean firing frequency of prospective PNs and fast spiking interneurons (FSI) during rest and movement, and the mean through-to peak time of spikes in control and ELS groups. PN firing rates: rest, control n=13 (5), ELS n=34(6), movement, control n=5(3), ELS n=17(4); p=0.087, F _(1, 65)_ = 3.027, 2-way ANOVA; *p=0.028, Mann-Whitney. FSI firing rates: rest, control n= 8(4), ELS n=16(6), movement control n= 4(3), ELS n= 8(3); p=0.13, F _(1, 32)_ = 2.42, 2-way ANOVA. PN through-to - peak time, control n=22(5), ELS n=35 (6), *p=0.013, Mann-Whitney; FSI through to peak time, control n=8(4), ELS n=14 (6), p=0.075, unpaired t-test.

Feedforward inhibition gates induction of synaptic plasticity and in particular, LTP induced by theta-burst stimulation (TBS) in the LA (Bissiere et al., 2003; Bazelot et al., 2015; Haikonen et al., 2022). To investigate whether the observed changes in the EPSC/dIPSC ratios in ELS exposed mice correlated with alterations in TBS-induced LTP, we recorded the postsynaptic field responses evoked by stimulation of the external capsulae in the LA. ELS treatment had no significant effect on TBS-induced LTP in males (Figure 5B). However, consistent with the strong feedforward inhibition, TBS-induced synaptic potentiation was smaller in ELS exposed females compared to controls (Figure 5C).

Thus, in adult ELS-exposed males, both the high intrinsic excitability of the LA principal neurons as well as weak feedforward inhibition shift the network towards hyperexcitability. In ELS females, the excitability of the LA principal neurons was only slightly elevated and is balanced by enhanced feedforward inhibition during activation of cortical inputs. However, induction of LTP was compromised selectively in ELS-exposed females.

### ELS is associated with LA hyperexcitability and disrupted circuit dynamics in adult male mice

To further investigate the LA circuit function, we performed *in vivo* recordings with the Neuropixels probes from LA in head-fixed mice, running on a treadmill. In these recordings we focused on males only, since their ELS-associated phenotypes related to both, circuit physiology as well as behavior were stronger than females in the adulthood. Even though the mice were carefully habituated to the recording conditions, the corticosterone levels during the head-fixation were high (head-fixed: 338 ± 21 ng/ml, control: 26 ± 4 ng/ml; p=0.0117, Mann-Whitney). Therefore, behaviorally, this recording configuration should be interpreted as repeated stress after the ELS.

To investigate firing patterns of individual neurons, the data was sorted into single units, which were further classified into prospective PNs and fast-spiking interneurons (FSIs) based on their spike waveform properties. The mean firing frequency of the putative LA PNs was higher in ELS-exposed mice than in controls when the mice were not moving (p=0.028, Mann-Whitney). No significant differences in the mean firing frequency of the FSIs were detected. Furthermore, in the ELS-exposed mice the PN spike waveforms, analyzed as the through-to-peak-time, were faster compared to controls (Figure 5D), consistent with LA hyperexcitability.

As aberrant patterns of oscillatory activity have been linked to stress and anxiety-like behaviors (Murthy and Gould, 2020; Antonoudiou et al., 2023) we went on to analyze the local field potentials in the LA from the same recordings. When the animals were not moving, the oscillatory power in control and ELS-exposed mice was similar, with slightly (but not significantly) elevated gamma power in the ELS group (Figure 6A,B). Theta power (4-12 Hz) was significantly increased in controls but not in ELS-exposed mice when the animals started moving. On the other hand, the power in mid-gamma frequency range (50-90 Hz) increased in ELS-exposed but not in control mice during transition to movement (Figure 6C). Thus, in ELS-exposed mice, the LA circuitry seems to be less dynamic in the theta frequencies and more dynamic in the gamma frequency range.

**Figure 6.**
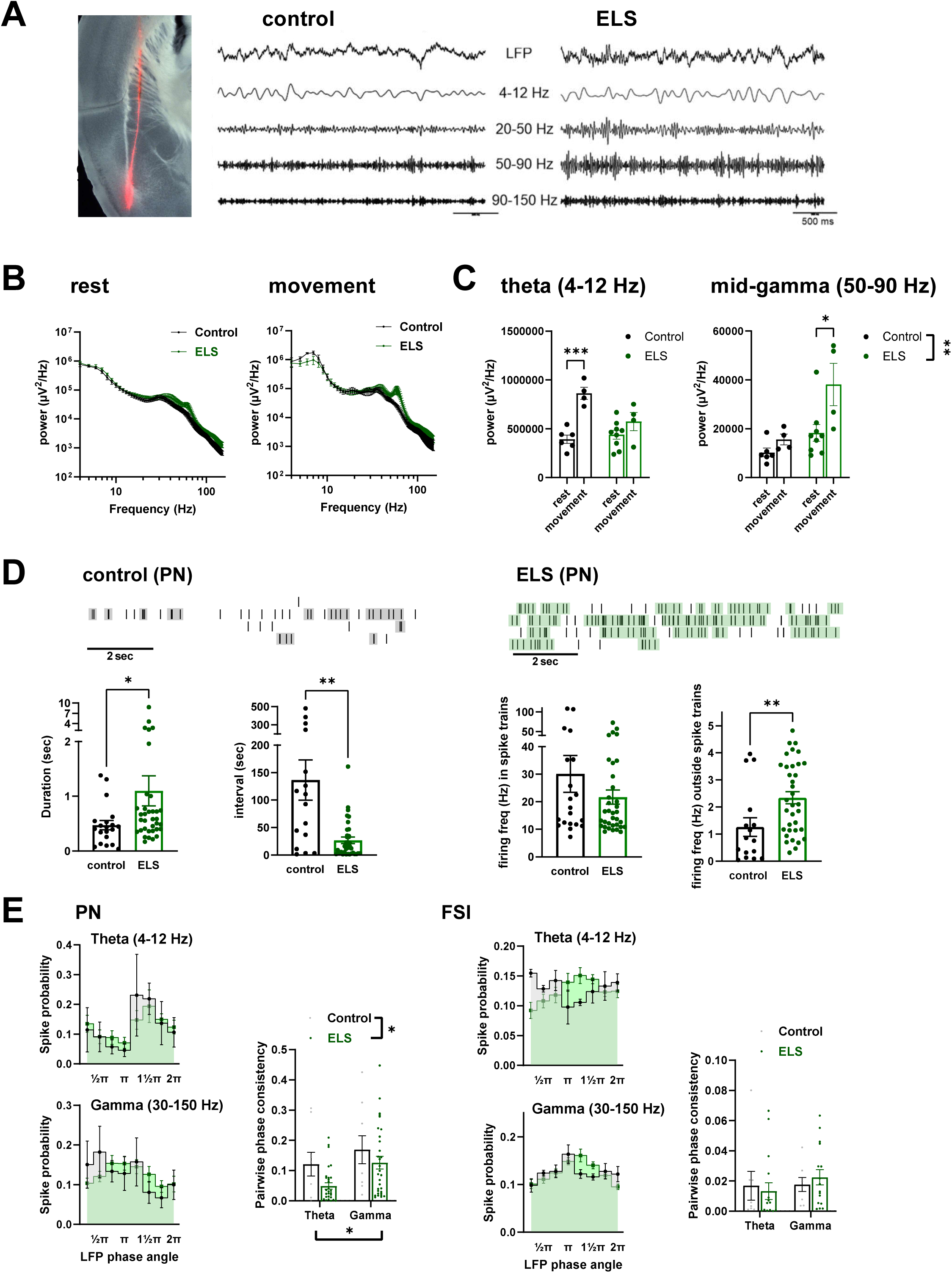
Oscillation dynamics in the LA of adult control and ELS-exposed mice. A. Image illustrating the localization of the fluorescently labelled recording electrode in a coronal section including the LA. The traces show an example of the LFP recording of oscillatory activity in the LA of control and ELS-exposed adult mice, and after band – pass filtering [theta (4-12 Hz), low (20-50 Hz), mid (50-90 Hz) and high (90-150 Hz) gamma]. B. LFP power spectra during rest, for control (black, n=6) and ELS exposed (green, n=9) mice, and during movement on the treadmill. For movement periods, n=4 / group, as some data was rejected because of movement artefacts. C. Oscillatory power in the theta (4 – 12 Hz) and mid-gamma (50-90 Hz) frequency bands for channels located in the LA, during rest and movement periods. For theta, effect of ELS, p=0.060, F_(1, 19)_ = 4.00, 2-way ANOVA; *** p= 0.0002, Holm-Sidak. For mid-gamma, effect of ELS, **p=0.0026, F _(1, 19)_ = 11.95, 2-way ANOVA; * p= 0.018, Holm-Sidak. D. Raster plots illustrating the firing patterns of single PN units in the LA of control and ELS exposed mice. The detected spike trains (bursts) are highlighted in grey (control) or green (ELS). Pooled data on the train duration (* p=0.023, Mann-Whitney), interval (***p=0.0017, Mann-Whitney) and firing frequency (p=0.439, Mann-Whitney) as well as on the firing frequency between the spike trains (**p=0.0062, Mann-Whitney). control, n=20, ELS, n= 35 units. E. Spike probability distribution across the phase of theta (4-12 Hz) and gamma (30-150 Hz) band LFP oscillation for phase locked PN and FSI units in the LA (PN, theta n= 5 and 12, gamma n= 9 and 32; FSI theta n=4 and 7, gamma n=6 and 13, for control and ELS, respectively). Analysis of the PPC (ELS effect in PN, * p=0.037; F_(1, 76)_ = 4.53; FSI, p=0.92, F_(1, 44)_ = 0.0095, 2-way ANOVA; frequency effect in PN, *p=0.023; F_(1,76)_ = 5.36).

As the LFP data suggested aberrant rhythmic activity in the LA of the ELS-exposed mice, we conducted more detailed analysis of the spiking patterns of individual neurons. This analysis was done for the rest periods only, given the low number of units detected during the movement epochs. Interestingly, LA PNs in the ELS-exposed mice were firing in irregular spike trains that were longer in duration and more frequent than in controls (Figure 6D). The firing frequency within these spike trains typically fell within the high theta - low gamma frequency bands (10-40Hz) and was not significantly different between the groups (Figure 6D). However, consistent with the hyperexcitability, the firing frequency of the PNs between the spike trains was higher in the ELS-exposed mice than controls (Figure 6D). In contrast to PNs, no significant differences in the spiking patterns were seen in FSIs (data not shown).

We also analyzed the relationship between single-unit firing patterns and LFP oscillations during the rest periods. In both control and ELS groups, most of the units representing PNs and FSIs were phase-locked to the gamma oscillations (PNs, control 78%, ELS 94%; FSIs, control 75%, ELS 81%), while a subpopulation of the units was locked to theta (PNs, control 33%, ELS 32 %; FSIs, control, 50%, ELS 44%). In PNs, analysis of pairwise phase consistency (PPC) indicated that entrainment of phase-locked units to gamma and theta was lower in ELS-exposed mice compared to controls (Figure 6E). We did not detect significant differences between the groups in the phase angle (F _(1, 45)_ = 0.375, p=0.54, 2-way ANOVA). In FSIs, no significant differences in the PPC (Figure 6E) or the phase angle (F _(1, 26)_ = 1.228, p=0.28, 2-way ANOVA) were detected between the groups. These data suggest that the high excitability of the PNs in the ELS-exposed mice results in increased burst firing, which however, is not fully synchronized to the population oscillations within the LA network.

## DISCUSSION

### Behavioral phenotype induced by ELS

MS and LBN represent the most investigated models of ELS. While MS mimics childhood neglect in humans (Lehmann and Feldon, 2000; Lupien et al., 2009), LBN represents a resource scarcity model with disrupted maternal care (Molet et al., 2014; Walker et al., 2017). Both models lead to aberrant emotional and cognitive behaviors in the adulthood, however, the exact outcome is highly dependent on the type of stress, developmental timing as well as sex (Bath, 2020; Ellis and Honeycutt, 2021; Waters and Gould 2022).

We chose to use a ‘double-hit’ model combining LBN with unpredictable epochs of MS, that has been previously reported to produce a robust male-specific anxiety phenotype that is associated with amygdala hyper-connectivity (Johnson et al., 2018). As shown before, rearing under these conditions resulted in disrupted care of the pups, evidenced by dam’s frequent nest exits (Molet et al., 2016; Walker et al., 2017). Additionally, pups from three different mouse strains exposed to the combined LBN + MS treatment gained less weight than controls, highlighting the reproducibility of the model across mouse lines (Molet et al., 2016; Walker et al., 2017). Consistent with previously observed defects in spatial learning and memory (Bath et al., 2017; Rocha et al., 2021), all ELS exposed mice showed impaired performance in contextual fear conditioning task as adults. However, only male mice showed elevated anxiety-like behavior in the OF test and impaired cued fear conditioning, which, compared to contextual conditioning, is thought to involve more exclusive role of the amygdala. In contrast to the previous work (Johnson et al., 2018), we did not observe anxiety behavior in the light-dark box or the elevated plus maze (EPM), which could reflect the shorter duration of the ELS in our protocol (P4-14 vs P0–25). OF is more sensitive for assessment of anxiety traits and the influences of environmental factors on anxiety-like behavior compared to the EPM (Goes et al., 2015; Cerqueira et al., 2023), which might be critical when the environmental exposure is less robust. Yet, the significant sex-specific defects in amygdala-associated behaviors provided an interesting model to investigate the effects of ELS on physiological functions of different cell types in the LA.

### Development of amygdala hyperexcitability after ELS

ELS result in amygdala hyperexcitability (Horii-Hayashi et al., 2013; Malter-Cohen et al., 2013; Raineki et al., 2019; Guadagno et al., 2020; Donati et al., 2024), yet very little is known about its physiological underpinnings at different stages of development. In adults, chronic stress increases the intrinsic excitability of the principal neurons in the LA (Rosenkrantz et al., 2010) and comparable findings have been obtained in the BLA after adolescent social isolation stress (Rau et al., 2015). In addition, chronic stress perturbs phasic and tonic inhibitory control of PNs in the adult BLA, which significantly contributes to their hyperexcitability (Ryazantseva et al., 2024; reviewed in Sharp 2017).

Our recordings revealed hyperexcitability of LA PNs soon after the ELS exposure in juvenile females, but not in males; in males a significant increase in excitability was only observed in the adult stage. As a notable contrast to adult stress, ELS was associated with developmentally transient increase in inhibitory tone in LA PNs of both sexes, involving elevated GABAergic inhibition (sIPSC frequency) and weaker excitatory inputs (sEPSC amplitude) in juveniles. In control animals the synaptic E/I balance was strongly downregulated during development, largely because of the developmental increase in GABAergic inhibition. Although sIPSC frequency in the PNs was only slightly elevated after the ELS treatment, this together with the weakened glutamatergic drive was sufficient to shift the E/I balance prematurely towards inhibition in ELS-exposed mice. Thus, in this respect, ELS was associated with accelerated development of the network, in both males and females.

The enhanced inhibitory tone during/after ELS could directly contribute to the delayed emergence of PN hyperexcitability in males and also trigger homeostatic adaptations in intrinsic excitability of the PNs to compensate for altered activity levels. Additionally, stress hormones may directly affect the excitability of LA PNs (Duvarci and Pare, 2007) and produce changes in gene expression that persist over time. Our data cannot explain why the PN hyperexcitability was observed in juvenile females but not in males; we can only speculate that the sex differences in timing of neurodevelopment and in epigenetic programming contribute to the observed differences (Bath, 2020).

### Effect of ELS on functional maturation PV and SOM interneurons

PV INs have specific electrophysiological and morphological properties that are essential for controlling the activity of the surrounding principal cells (Hu et al., 2014). Their fast-spiking characteristics develop gradually during the first postnatal weeks of life in rodents (Doischer et al., 2008; Goldberg et al., 2011), coinciding with the time of typical ELS exposure. Previous work has suggested that LBN facilitates neurochemical maturation of PV INs in the amygdala (Manzano Nieves et al., 2020; Guadagno et al., 2020), while MS typically leads to a reduction in PV intensity in the amygdala (Lukkes et al., 2018; Gildawie et al., 2020; Soares et al., 2020; Haikonen et al., 2022). Precocious maturation of functional and morphological properties of PV INs could readily explain the elevated GABAergic tone that was observed in juvenile ELS-exposed mice. However, we did not detect significant ELS-dependent alterations in the density of PV or SOM cells in the juvenile LA in our ‘double-hit’ model that combined LBN and MS. Nonetheless, electrophysiological recordings revealed substantial ELS-related changes in the functional properties of these INs.

First, ELS was associated with a developmentally transient increase in glutamatergic synaptic inputs to GABAergic interneurons in the LA. Elevated glutamatergic input was observed in PV and SOM interneurons in both juvenile males and females after ELS exposure. It is plausible that hyperexcitability of the PNs in ELS females contributes to the excitatory drive to at least SOM INs, where the effect of ELS on sEPSCs was most pronounced. In addition, we have previously shown that MS increases projections from mPFC to amygdala GABAergic neurons in male rats (Haikonen et al., 2022), and corresponding mechanism involving ELS-dependent modulation of long-range excitatory inputs to LA INs may be assumed here as well.

Second, the excitability of PV and SOM INs in ELS juveniles was lower compared to controls and was associated with low input resistance. Similar phenotype was recently described in LA PV INs after repeated restraint stress in adults (Ryazantseva et al., 2024), suggesting that the K+ channels that regulate the rheobase and input resistance of amygdala PV INs are modulated by chronic stress independently of the developmental stage. In ELS-exposed mice, this effect was developmentally transient and normalized towards adulthood, where the excitability of the INs in the ELS group was comparable to controls or even slightly elevated (SOM INs).

Overall, the effects of ELS on the juvenile PV and SOM INs were remarkably similar between males and females. However, during the development to adulthood, differences between the sexes emerged: while glutamatergic inputs to GABAergic neurons developed prematurely in both ELS-exposed males and females, there was no further increase in these inputs in males after weaning, resulting in low connectivity in the adult stage. ELS females seemed to resist this pathological trajectory and in adults, the overall glutamatergic drive to interneurons was comparable to controls. In both sexes, the firing properties of PV and SOM were normalized in adulthood following the transient perturbation after the ELS exposure.

As the amygdala-dependent behaviors and fear processing were more severely affected in males, these results are consistent with the idea that perturbed or arrested development of glutamatergic synaptic inputs to GABAergic interneurons after the ELS exposure correlates with the emergence of pathological behavioral phenotypes in male mice.

### LA microcircuit function in ELS exposed adults

PV and SOM INs are critical components of the feedforward inhibitory circuits that gate and pace the activity of principal neurons in the LA in response to afferent activation (Krabbe et al., 2018). Consistent with impaired glutamatergic connectivity of these INs, feedforward inhibition in response to activation of cortical afferents was significantly weaker in ELS males than in controls. In contrast, the strength of feedforward inhibition was elevated in ELS females, likely because of the higher excitability of SOM INs that became decisive in the absence of differences in the strength of excitatory inputs.

GABAergic transmission has a pivotal role in gating plasticity of afferent inputs in LA (Bissiere et al., 2003; Shin et al., 2006; Haikonen et al., 2022). Accordingly, the elevated GABAergic drive was associated with attenuated LTP in the cortical afferents in ELS-exposed females. Impaired LTP has been previously reported in the LA after MS (Haikonen et al., 2022; Danielewic and Hess, 2014), and in the hippocampus of LBN-treated mice (Lesuis et al., 2019), where it correlates with aberrant fear memory. Here, attenuated LTP was observed only in females, but fear-related memory was impaired in both sexes, suggesting that also other factors contribute to the cognitive defects observed in ELS-exposed adult mice.

### Effect of ELS on LA excitability and oscillatory dynamics in adult males in vivo

The *in vitro* electrophysiological data indicate that in adult ELS-exposed males, both the high intrinsic excitability of the LA principal neurons as well as weak feedforward inhibition shift the network towards hyperexcitability. Consistently, the firing rate of putative LA PNs was elevated in ELS-exposed mice *in vivo*. The resting membrane potential of LA PNs in ELS group was slightly depolarized compared to the controls. This depolarization could facilitate burst firing, as BLA neurons have an inherent tendency to produce voltage-dependent membrane potential oscillations (Pape and Pare, 2010). Accordingly, the firing pattern of LA PNs in ELS exposed mice was characterized by irregular spike trains that were longer and more frequent compared to controls. Furthermore, firing of individual PNs in the ELS group was less well entrained to the population synchrony than in controls, consistent with the idea that ELS facilitates uncoherent firing of the PNs.

Dynamic alterations in both theta and gamma oscillations in the amygdala have been linked to various behaviors, including fear, stress, and anxiety (Murthy and Gould, 2020; Antonoudiou et al., 2023). Specifically, chronic stress has been shown to enhance the gamma oscillation power in the amygdala in response to behavioral stimuli, including auditory stimulus in the adults (Ghosh et al., 2013), and social interaction in the juveniles (Raineki et al., 2019). In our model, head fixation induced acute stress in the recorded mice, which could affect network activity in the LA. Although

ELS results in enhanced sensitivity to stress later in life (Pena et al., 2017), we detected no differences between ELS and control mice in the LFP power spectra when the animals were at rest. However, when the animals started moving on the treadmill, gamma power was significantly elevated in the LA of ELS-exposed mice compared to controls. In parallel, dynamic changes in the theta power were reduced.

Gamma oscillations efficiently synchronize local circuits in the amygdala during memory consolidation (Amir et al., 2018; Headley et al., 2021). The persistent enhancement of gamma oscillation dynamics in the LA after ELS exposure might influence encoding of emotionally salient memories. Additionally, gamma activity in the BLA has been specifically linked to a state of apprehension (Amir et al., 2018; Headley et al., 2021), correlating enhanced gamma to fear generalization after ELS.

Theta oscillations are thought mediate amygdala interactions with mPFC and ventral hippocampus during discrimination of fear and safety and expression of anxiety-like behaviors (Likhtik et al., 2014; Murthy and Gould, 2020; Pare and Headley 2023). Hence, loss of the theta oscillation dynamics in ELS exposed mice is consistent with impaired functional connectivity between corticolimbic brain regions, which has been repeatedly observed after ELS (Reincke and Hanganu-Opatz 2017; Guadagno 2018; Honeycutt 2020; Haikonen et al., 2022; Donati et al., 2024).

Interestingly, glutamatergic inputs to interneurons play a pivotal role in regulating gamma oscillations within cortical and limbic circuits (Korotkova et al., 2010; Carlen et al., 2012; Ojanen et al., 2023; Hadler et al., 2024) and have also been implicated in long-range theta synchronization in the cortico-limbic circuits (e.g. Bazelot et al., 2015; Yang et al., 2021). As ELS had no significant effects on other physiological properties of PV and SOM INs in adult males, our data is consistent with the idea that the perturbed connectivity of interneurons is a key mechanism underlying the aberrant oscillatory dynamics in the LA in ELS exposed male mice. Imbalance in the oscillatory states in the amygdala and across brain regions after ELS may contribute to the enhanced emotional symptoms in stress-related psychiatric disorders (Murthy and Gould, 2020; Antonoudiou et al., 2023; Pare and Headley 2023).

## Author contributions

V.S. setting up the ELS model, in vitro electrophysiology (experiments and data analysis), immunostainings and behavioral testing

A.L., H.H. setting up the ELS model, analysis of the behavioral data

J.H. in vitro electrophysiology, LTP recordings (experiments and data analysis)

A-J.K. immunohistological stainings, in vitro electrophysiology (experiments and data analysis)

M.P. in vitro electrophysiology

J-K.R., S.O. in vivo electrophysiology and data analysis

O.L. analysis of the in vivo electrophysiological data

S.E.L. Conceptualization and supervision of the project, funding acquisition

S.E.L. wrote the paper with input from all other authors.

## Acknowledgements

We thank the personnel and facilities provided by the following HiLife research infrastructures: Mouse Behavioral Phenotyping Facility, Rodent In vivo Brain Imaging and Electrophysiology unit and the Laboratory Animal Center. Janne Sulku is acknowledged for expert technical help. We wish to acknowledge CSC – IT Center for Science, Finland, for computational resources. This study was financially supported by the Academy of Finland, Sigrid Juselius Foundation, Jane and Aatos Erkko foundation and Finnish Cultural Foundation. V.S. was supported by Orion Research Foundation. Mouse Behavioral Phenotyping Facility and Helsinki Electrophysiology Platform are supported by Biocenter Finland.

---

**Supplementary Figure 1.**
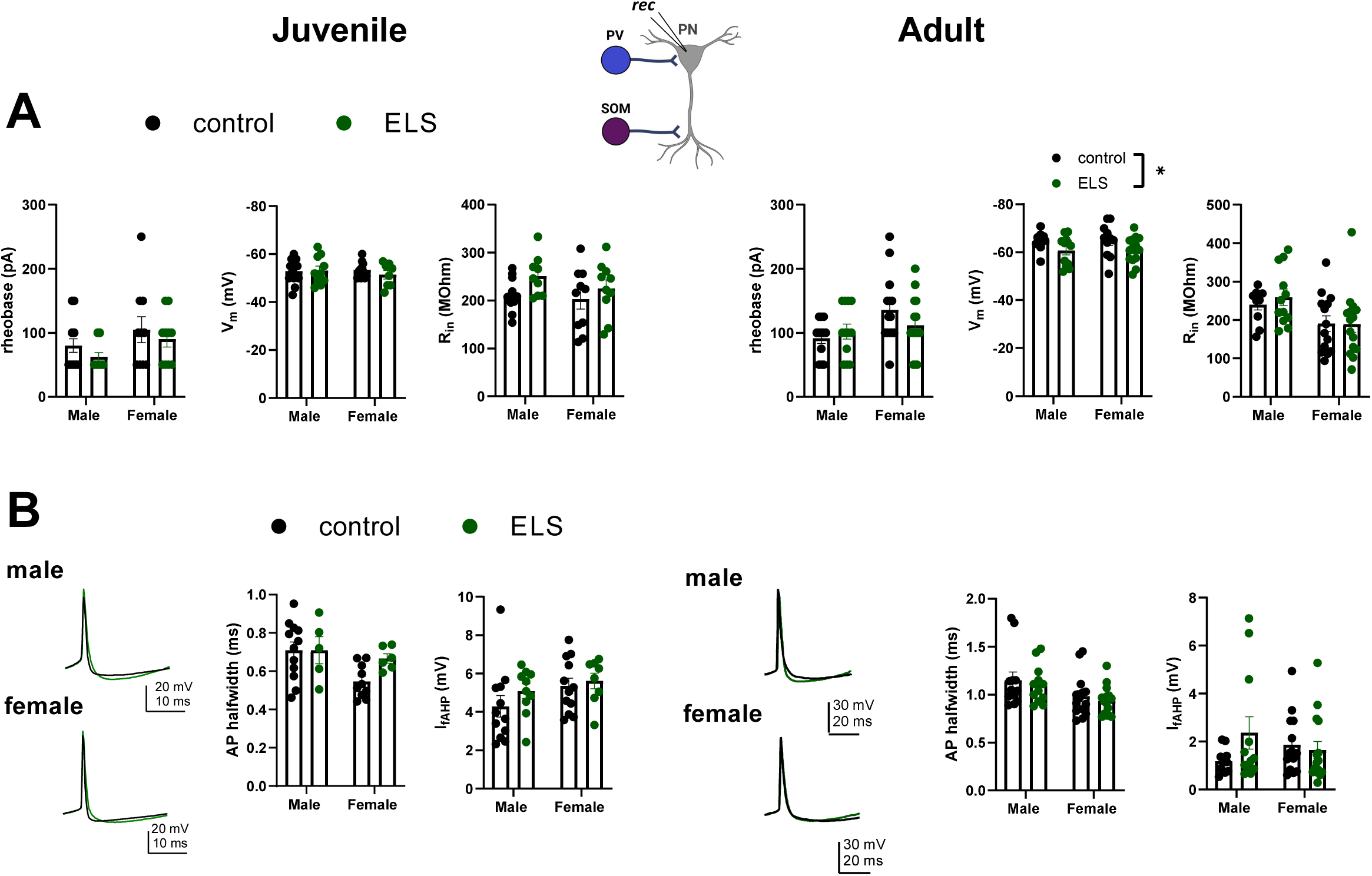
Membrane properties of the LA PNs for the recordings shown in 2A and 2B. A. Pooled data on the rheobase, resting membrane potential (Vm) and input resistance (R_in_) in control and ELS exposed males and females, at the two developmental stages. No significant differences between the groups were detected in juveniles. In the adults, ELS had a significant effect on the resting membrane potential (F _(1, 43)_ = 5,628; *p=0.022, 2-way ANOVA). B. Expanded example traces and pooled data on the AP waveform control (black) and ELS exposed (green) males and females, at the two developmental stages. ELS treatment had no significant effects on the AP kinetics in either group.

**Supplementary Figure 2.**
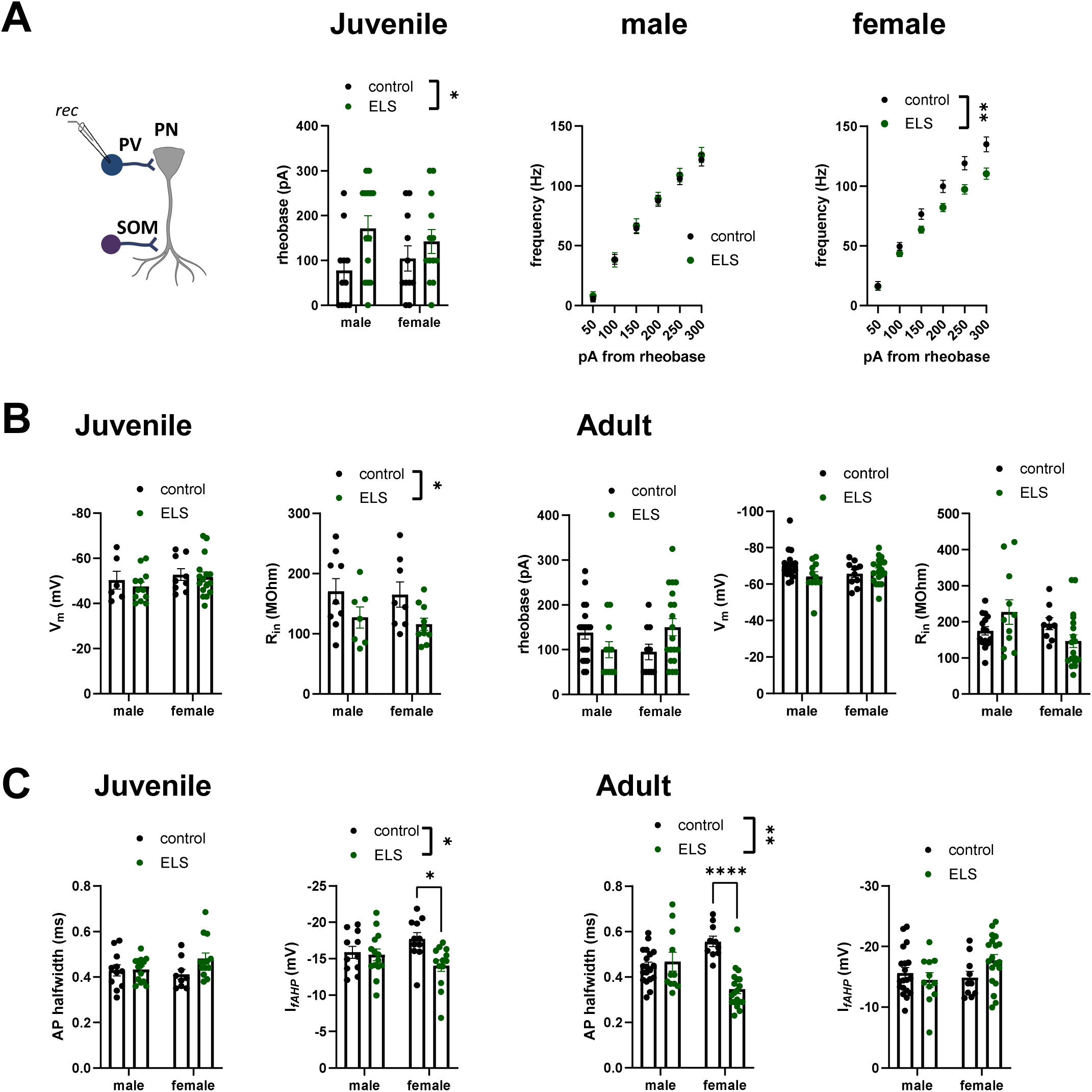
Membrane properties of the PV INs for the recordings shown in 3A and 3B. A. Pooled data on the rheobase in PV INs in juvenile control and ELS exposed mice (left), and the PV INs firing rate, for the same data as in Figure 3A, normalized to the rheobase. Effect of ELS on rheobase F_(1, 45)_ = 5.758, *p=0.021, 2-way ANOVA. Effect of ELS on rheobase-adjusted firing rate in males, F_(1, 23)_ = 0.1207, p=0.73; females, F_(1, 21)_ = 9.606, **p= 0.0054, RM ANOVA. B. Pooled data on the resting membrane potential (Vm) and input resistance (R_in_) in control and ELS exposed males and females, at the two developmental stages, and for rheobase in adults. ELS had a significant effect on input resistance (F_(1, 30)_ = 7.005, *p=0.0128) in juveniles, but not in adults . C. Pooled data on the effect of ELS on AP waveform. ELS treatment has a significant effect on the amplitude of the fast AHP current in juveniles (F _(1, 45)_ = 5.901, *p=0.0192, 2-way ANOVA: * p=0.0166, Tukey’s test for females) and on AP halfwidth in adults (F _(1, 55)_ = 14.04, **p=0.004, 2-way ANOVA, **** p<0.0001, Tukey’s test for females).

**Supplementary Figure 3.**
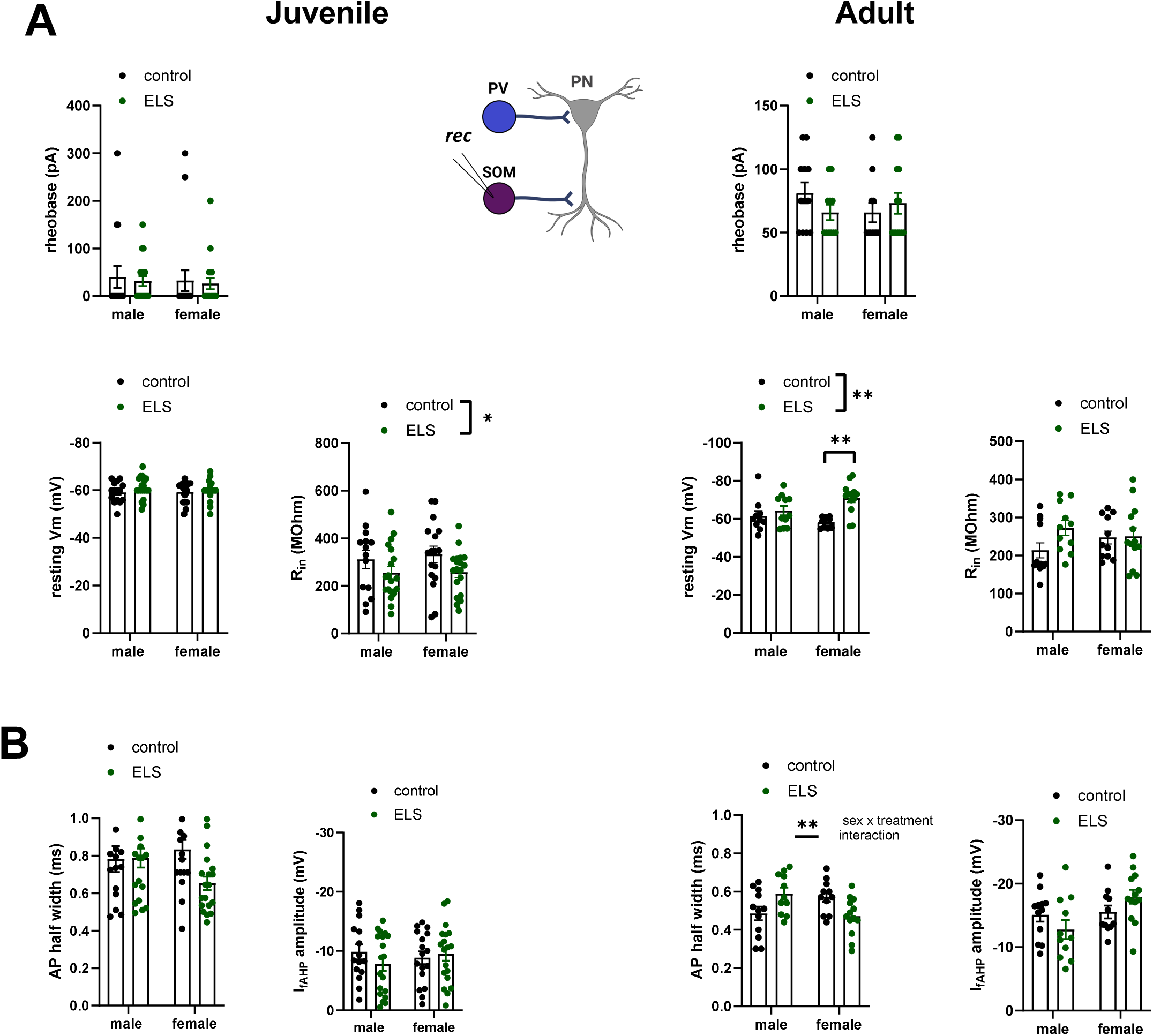
Membrane properties of the SOM INs for the recordings shown in 4A and 4B. A. Pooled data on the rheobase, resting membrane potential (Vm) and input resistance (R_in_) in control and ELS exposed males and females, at the two developmental stages. ELS had a significant effect on the input resistance in the juveniles (F_(1, 65)_ = 4.976, *p=0.0292) and on resting membrane potential in the adults (F_(1, 40)_ = 12.17, **p=0.0012; **p= 0.0011 Tukey’s test). B. Pooled data on the effect of ELS on AP kinetics at the two developmental stages. ELS treatment has a significant sex-dependent effect on the AP halfwidth in the adults (interaction, F_(1, 43)_ = 11.37, **p=0.0016, 2-way ANOVA).

## METHODS

### Animals

Experiments were performed in male and female mice at two developmental time points (juveniles: P18-21, adults: P60-90). The following mouse lines were used : WT C57BL/6J (JAX 000664), Grik1^tm1c/tm1c^ homozygotes (described in Ryazantseva et al., 2020). PV-Cre (JAX 008069) and Sst-ires-Cre (Sst^tm2.1(cre)Zjh/ J^ , kindly provided by Dr. Juzoh Umemori) lines were crossed with Ai14 tdTomato reporter (JAX 007914). All genetically modified mice were of C57BL/6J background. The mice were group housed (2–5 animals of the same sex/cage) in a temperature and humidity-controlled environment (22±2 °C; 50±15 %), with a 12/12h light/dark cycle (lights on 6 am., off 6 pm) and *ad libitum* access to food and water. The cages were enriched by nesting material (aspen strips, PM90L, Tapvei) and aspen bricks (50 × 10 × 10 mm, Tapvei) and handling tube (red or transparent, length 100 mm, diameter 45 mm). The animals were sacrificed for experiments between 8 and 10 a.m., during the light period. The stage of the estrous cycle was not controlled. All the experiments were done in accordance with the University of Helsinki Animal Welfare Guidelines and approved by the Animal Experiment Board in Finland (license numbers: KEK-17-019, KEK22-010, ESAVI/29384/2019, and ESAVI/31984/2022).

### Early life stress

Early life stress was induced with limited bedding and nesting (LBN) model, combined with maternal separation (MS). Only females that previously gave birth were subjected to ELS to reduce the risk of pup abandonment and filial cannibalism. Only litters with at least 4 pups but no more than 8 pups and at least one male and one female pup in the litter were used for both control and LBN animals. For litters of larger sizes additional pups were sacrificed at P4. LBN litters pups were kept on a raised wire mesh platform with no bedding and half a Nestlet (2.5 g compressed cotton square; Ancare, Bellmore, NY) for nesting material (Walker et al., 2017). LBN conditions lasted from P4 until P14. Additionally, we included maternal separations at P8, P10 and P12 as an unpredictable stressor, which has been shown to produce a more robust anxiety phenotype (Johnson et al., 2018). The 1h long separation was performed between 9 am and 12 am by moving the pups into a separate warmed cage with heat pads in the same room. All the pups were kept together during the separation period. The pups were weighed at P4 and P14 and their individual weights were recorded by sex. Control animals of matching genotype were raised in normal conditions and the pups were also weighed at P4 and P14.

### Behavioral tests

#### Maternal behavior

was assessed at P5 and P10 at two time points – between 10 and 11 a.m. (light part of the cycle) and between 7 and 8 p.m. (dark part of the cycle). The room was set up with surveillance cameras recording to a local PC to mitigate any additional disturbances to the animals. The footage was then manually evaluated. The dam was counted as “out” of the nest when none of its paws, head or body were in contact with the main group of the pups.

#### Elevated plus-maze (EPM)

The maze consisted of two open arms (30 x 5 cm) and two enclosed arms (30 x 5 cm) connected by central platform (5 x 5 cm) and elevated to 40 cm above the floor. Open arms were surrounded by 0.5 cm high edges to prevent accidental falling of mice. The floor of each arm was light grey and the closed arms had transparent (15 cm high) side- and end-walls. The illumination level in all arms was ∼150 lx. The animal was placed in the center of the maze facing one of the closed arms and observed for 5 minutes. The trials were recorded with Ethovision XT13 videotracking software (Noldus Information Technology, Wageningen, The Netherlands). The latency to the first open arm entry, number of open and closed arm entries (four paw criterion) and the time spent in different zones of the maze were measured. The number of faecal boli was counted after trial.

#### Light-dark box (LD)

The test was carried out in the open field arena (30 x 30 cm, Med Associates, St. Albans, VT) equipped with infrared light sensors detecting horizontal and vertical activity. The dark insert (non-transparent for visible light) was used to divide the arena into two halves, an opening (a door with a width of 5.5 cm and height of 7 cm) in the wall of the insert allowed animal’s free movement from one compartment to another. Illumination in the center of the light compartment was ∼550 lx (bright ceiling lights). Animal was placed in the dark compartment and allowed to explore the arena for 10 minutes. Distance travelled, number of rearings, latency to enter the light compartment and time spent in different compartments were recorded by the program. The number of faecal boli was counted by experimenter after the end of trial.

#### Open field (OF)

The mice were released in the corner of novel open field arena (white floor, transparent walls, 30 x 30 cm, Med Associates). Horizontal and vertical activity (distance moved and rearings, respectively) was recorded by infrared sensors for 30 minutes (light intensity ∼150 lx). Peripheral zone was defined as a 6 cm wide corridor along the wall.

#### Fear conditioning (FC)

The experiments were carried out as described (Haikonen et al., 2024), using a computer-controlled fear conditioning system (TSE Systems, Germany). Training was performed in a transparent acrylic cage (23 × 23 × 35 cm) within a constantly illuminated (∼ 100 lx) fear conditioning box. A loudspeaker provided a constant, white background noise (68 dB) for 120 s followed by 10 kHz tone (CS, 76 dB, pulsed 5 Hz) for 30 s. The tone was terminated by a footshock (US, 0.6 mA, 2 s, constant current) delivered through a stainless steel grid-floor (bar Ø 4 mm, distance between the bars 10 mm). Two CS-US pairings were separated by a 30 s pause. Contextual memory was tested 24 h after the training. The animals were returned to the conditioning box and total time of freezing (defined as an absence of any movements for more than 2 s) was measured through infrared light beams scanned continuously with a frequency of 10 Hz. The CS was not used during this time. Memory for the CS (tone) was tested 2 h later in a novel context. The new context was a similarly sized acrylic box with black non-transparent walls and smooth floor. A layer of wood chips (clean bedding material) under the floor provided a novel odour to the chamber. After 120 s of free exploration in a novel context the CS was applied for additional 120 s and freezing was measured as above.

### In vitro Electrophysiology

#### Acute slices

For adult animals (P60-P80), acute coronal sections were prepared as previously described (Englund et al., 2021). Briefly, the mice were anesthetized with isoflurane until the loss of pinch reaction. The animal was then quickly decapitated and brain was removed and immediately placed in carbogenated (95% O_2_ / 5% CO_2_ ) ice-cold N-Methyl-D-glucamine (NMDG) based solution (pH 7.3–7.4) containing (in mM): 92 NMDG, 2.5 KCl, 1.25 NaH_2_PO_4_, 30 NaHCO_3_, 20 HEPES, 25 glucose, 2 thiourea, 5 Na-ascorbate, 3 Na-pyruvate, 0.5 CaCl_2_ and 10 MgSO_4_. The vibratome (Leica VT 1200S) was used to obtain 300 µm thick brain slices. Slices containing the amygdala were placed into a slice holder and incubated for 8–10 min at 34°C in the NMDG–based solution. Slices were then transferred into a separate slice holder at room temperature with a solution containing (in mM): 92 NaCl, 2.5 KCl, 1.25 NaH_2_PO_4_, 30 NaHCO_3_, 20 HEPES, 25 glucose, 2 thiourea, 5 Na-ascorbate, 3 Na-pyruvate, 2 CaCl_2_ and 2 MgSO_4_ (continuously perfused with 95% O2 / 5% CO2). For juvenile animals (P18-P21), similar method was used, but the dissection solution contained (in mM):124 NaCl, 3 KCl, 1.25 NaH_2_PO_4_, 10 MgSO_4_, 26 NaHCO_3_, 15 glucose, 1 CaCl_2_, ice-cold and saturated with 5% CO_2_/95% O_2_. The slices were transferred to a slice holder containing ACSF with elevated MgSO_4_ concentration (4mM MgSO_4_) and incubated for 30 min at 34°C. Then the slices were kept for another 30 min at room temperature before recording.

#### Patch-clamp recordings

Whole-cell patch-clamp recordings were done from LA neurons using standard methods (Lauri et al., 2006). After 1–4 h of recovery, the slices were placed in a submerged heated (30-32 °C) recording chamber and continuously perfused with standard ACSF, containing (in mM): 124 NaCl, 3 KCl, 1.25 NaH_2_PO_4_, 26 NaHCO_3_, 15 glucose, 1 MgSO_4_ and 2 CaCl_2_, at the speed of 1.5–2 ml/min. Whole-cell patch-clamp recordings were done from lateral amygdala neurons under visual guidance using glass capillary microelectrodes with resistance of 3–5.5 MΩ. Multiclamp 700B amplifier (Molecular Devices), Digidata 1322 (Molecular Devices) or NI USB-6341 A/D board (National Instruments), and WinLTP version 2.20 (Anderson and Collingridge, 2007) or pClamp 11.0 software were used for data collection, with low pass filter (10 kHz) and a sampling rate of 20 kHz. In voltage-clamp recordings, uncompensated series resistance (Rs) was monitored by measuring the peak amplitude of the fast whole-cell capacitance current in response to a 5 mV step. Only experiments where Rs <30MΩ, and with <20% change in Rs during the experiment, were included in the analysis.

Lateral amygdala principal neurons (PNs) were initially selected based on cell size and morphology, then the cell type was confirmed based on the neuronal firing properties (firing frequency, spike frequency adaptation, AP half-width). Interneurons were identified with fluorescent markers.

#### Intrinsic excitability and spontaneous synaptic currents

Whole-cell current clamp recordings of membrane excitability were performed at resting membrane potential. Microelectrodes were filled with a solution containing (in mM): 135 K-gluconate, 10 HEPES, 5 EGTA, 2 KCl, 2 Ca(OH)_2_, 4 Mg-ATP, 0.5 Na-GTP (280 mOsm, pH 7.2). Depolarizing current steps of 400 ms duration were applied to induce action potential firing. The amplitude of the injected current was increased in 50 pA increments. Spontaneous excitatory and inhibitory synaptic currents (sEPSCs and sIPSCs) were recorded in whole-cell voltage-clamp at -70 mV and 0 mV respectively, using the same filling solution.

#### Evoked responses

EPSCs were recorded from principal neurons in LA at a holding potential of −70 mV using a cesium-based electrode filling solution (pH 7.2, 280 mOsm) containing (in mM): 125.5 Cs-methanesulfonate, 10 HEPES, 5 EGTA, 8 NaCl, 5 QX314, 4 Mg-ATP, 0.3 Na-GTP. EPSCs were evoked with bipolar stimulation electrodes (nickel-chromium wire), placed at external capsulae to mimic the activation of cortical afferents. Holding potential was then changed to 0 mV and dIPSCs were recorded. To confirm the disynaptic nature of the response CNQX 10 mM was applied at the end of the recording. Only recordings showing 80% or more inhibition were used for the analysis.

#### Field LTP recordings

LTP recordings from the LA were performed essentially as described previously (Haikonen et al., 2022). In order to obtain stable recordings, slices were allowed to sit in the recording chamber for 15 – 30 minutes before the experiment was started. fEPSPs were evoked with a bipolar stimulation electrodes (nickel-chromium wire) placed in the external capsule to mimic activation of cortical inputs, while the recording electrode (resistance 1-3 MΩ, filled with standard ACSF) was placed in the lateral amygdala (LA). The stimulation intensity was adjusted to produce a half-maximal response size. After a stable baseline, LTP was induced using theta-burst stimulation (6 bursts of 8/100Hz at 5Hz), repeated 3 times with 5 min intervals. WinLTP program was used for data acquisition (Anderson and Collingridge, 2007).

#### Data analysis

sIPSCs and sEPSCs were identified in Clampfit 10.0 software (Molecular Devices) using template search algorithm. The average amplitude and frequency of the detected events were calculated from at least 200 events per cell. Action potential frequencies were analyzed using the threshold search algorithm in Clampfit software. AP properties were analyzed from the 3rd spike in the train. Rin was calculated based on the response to -50 pA step under current clamp.

For EPSC/dIPSC ratios, 7-10 responses were averaged in each experimental condition and the peak values were assessed in Clampfit. EPSC/dIPSC ratio was calculated as the amplitude ratio of response at -70 mV/response at 0 mV.

In LTP experiments, WinLTP program was used to calculate the peak amplitude and slope of the evoked synaptic responses. For averaged data, the responses were normalized to the baseline. Mean potentiation was by comparing the averaged fEPSP slope calculated over a 5 minute period before and 60 min after the TBS.

For all the electrophysiological data, n number refers to the number of recorded neurons, which in all datasets were collected from at least three different animals. The number of animals used in each dataset is indicated in brackets after the n value.

### Statistical analysis

Statistical analysis was carried out on raw (not normalized) data using Graph Pad Prism 8.0.2 software. The sample size was based on previous experience on similar experiments. Normality of the distribution was checked with Shapiro-Wilk test, followed by two-way ANOVA with Holm-Sidak or Tukey’s post-hoc comparison or RM ANOVA, as appropriate. The results were considered significant at p<0.05. All the pooled data are given as mean ± S.E.M.

### Immunohistological staining

The mice were transcardially perfused with PBS, followed by 4% PFA under deep ketamine+xylazine anesthesia. The brains were extracted and postfixed at 4% PFA overnight (+4°C), after which the brains were stored in +4°C PBS before slicing the brain to 40 µm thick slices in PBS using a vibratome (Leica VT 1200S). Slices were incubated in blocking solution for 1 hour at room temperature (10% normal goat serum, 0,5% TritonX-100 and 1% BSA in PBS) and washed for 20 minutes with washing buffer (1% normal goat serum, 0,5% TritonX-100 and 1% BSA in PBS). Slices were then incubated overnight at +4°C with primary antibodies diluted in washing buffer (Anti-Parvalbumin Mouse SWANT PV235 1:1000, Anti-Somatostatin Rabbit Sigma-Aldrich SAB4502861 1:1000). After the overnight incubation, slices were washed 2 x 20 minutes with washing buffer and incubated with secondary antibodies in washing buffer for 1 hour at room temperature (Goat anti-mouse Alexa Fluor 488 1:1000, Goat anti-rabbit Alexa Fluor 568 1:1000), followed by 3 x 20 minutes of PBS washes. Slices were mounted with Vectashield mounting medium and sealed with clear nail polish.

The LA was imaged with Olympus BX63 fluorescent microscope with Olympus DP74 camera using the UPlanSApo x10/0.40 ∞/0.17/FN26.5 objective and Dimensions CellSens (3.2) software. Images were further analyzed with ImageJ software. Cell numbers within the LA were manually counted in a hand drawn area (in mm²) on 10x magnification images. The mean fluorescence intensity of PV and SOM neurons in the LA was measured from gray scale (8-bit) images with subtracted background.

### Corticosterone ELISA

Immediately after the last training session or the recording session, the mouse was placed under an upside down fuzzy fabric lined glass beaker that obstructed its view of the outer environment. About 50 μl of the tail blood was collected using an unrestrained tail snip method (Kim 2018) using a razor blade, a heparin coated pipette tip, and a heparin coated 200 μl centrifuge tube. Blood collection was done within 3-5 mins from the time of tail snip to minimize the confounding stress response; corticosterone level increase from the stress of blood sampling was reported to be not significant when done within 5 mins (Kim 2018). Tail was cauterized to stop the bleeding. Blood was centrifuged at room temperature for 10 mins at 12000 rpm (MC-12Pro, Joan Lab, Huzhou, China) to separate the plasma from the red blood cells, and at least 20 μl of plasma was transferred to a new 200 μl centrifuge tube and placed in a -80 °C freezer for further processing. Plasma corticosterone concentration was measured in duplicate using an enzyme-linked immunosorbent assay (ADI-900-097 corticosterone ELISA kit, Enzo life sciences, USA) and a microplate reader (Multiskan FC, Thermo Scientific, Waltham, MA, USA). The raw absorbance data was analyzed in Excel (Microsoft, Redmond, WA, USA).

### In vivo electrophysiology

#### Head plate implantation surgery

Head-plate implantation and craniotomies were performed at P80-90 essentially as described (Ojanen et al., 2023). Male mice (P80-90) were anesthetized using ≥ 4% isoflurane in an induction chamber. Carprofen (Rimadyl vet, 5 mg/kg s.c.), dexamethasone (2 mg/kg s.c.) and buprenorphine (Bupaq vet, 0,05 mg/kg s.c.) were injected, and the mice were placed on a 37 °C heating pad in the stereotaxic device. The depth of anesthesia was adjusted to 1,5 – 2 % isoflurane and both eyes were covered with ophthalmic ointment (Viscotears, Novartis). The hair on top of the head was trimmed and the skin was disinfected with povidone-iodine (Betadine). Lidocaine (0,5%, max 5 mg/kg) was injected under the scalp for local anesthesia and scalp and periosteum were removed. The surface of the skull was roughened using a large diameter drill bit and carefully cleaned. Tissue adhesive (Vetbond, 3M) was used to seal the edges of the wound. The locations for craniotomies were identified and marked, and small circular craniotomies were drilled (ML: -3,17 mm, RC: -1,6 mm for BLA and ML: -3,3 mm, RC: -3,3 mm for hippocampus). For a separate experiment, 300 nl of AAV [mDlx-EGFP] was injected to three injection sites in the hippocampus at 60 nl/min with 3 min waiting time between injections, using a glass capillary pipette. The head plate (model 5, Neurotar, Helsinki, Finland) was attached to the skull using a small amount of glue (Loctite precision), and a grounding screw was attached to the skull using dental cement (RelyX, 3M). The sides of the head plate and all visible areas of the skull were covered with dental cement and cured with UV light. The craniotomy sites were covered with silicone adhesive (Kwik-Sil, World Precision Instruments). The animals were allowed to recover on the heating pad, and when awake, placed in the home cage with some water-soaked soft food pellets. The weight of the mice was monitored and carprofen (Rimadyl vet, 5 mg/kg s.c.), and buprenorphine (Bupaq vet, 0,05 mg/kg s.c.) were injected for two days after the surgery for post-operative care. The mice were allowed to recover in the home cage for at least one week before starting the habituation protocol.

#### Habituation, craniotomy and electrophysiological recording from awake, head-fixed mice

The mice were habituated to being head-fixed on treadmill (SpeedBelt, Phenosys, Berlin, Germany) for 5 minutes, 10 minutes, 15 minutes, 20 minutes, 30 minutes, 45 minutes and 1 hour on consecutive days before the recordings. For recording, a bath was constructed on top of the opening of the head plate by making approximately 10 mm high walls out of Kwik-Sil, and the Kwik-Sil covering the craniotomies was removed. The ground electrode was attached to the grounding screw using silver wire, and the probe (Neuropixels 1.0) was stained by a DiI (V22885, TermoFisher) droplet, to allow post-experimental histological evaluation. The probe was placed on the surface of the brain in the middle of the predrilled craniotomy (ML: -3,17 mm, RC: -1,6 mm) and inserted horizontally to the depth 3,9 – 4,3 mm from dura mater. The bath on top of the head plate was filled with sterile filtered saline (0.9% NaCl). After 10 minutes stabilization period, simultaneous recordings of local field potential (LFP) and multi-unit activity (MUA) were performed at 30 kHz sampling rate (2500 Hz online down sampling for the LFP signal) using the IMEC Neuropixels acquisition system and Spike GLX software. After the recording, the craniotomy was covered with Kwik-Sil and the mouse was returned to the home cage. Within one week from recording the mice were transcardially perfused with PBS followed by 4% PFA and the brains were collected for histological verification of the electrode position.

### In vivo data analysis

#### Detection of epochs of rest and movement

The analog voltage output from the SpeedBelt was recorded simultaneously with the electrophysiological signals using the Spike GLX software, and voltages crossing the preset threshold of 10 mV from baseline were considered movements. Epochs of minimum 3 seconds of non-movement immediately after movement were included as non-movement epochs, and all continuous periods of movement were split into 3 second epochs and included in the analysis. All epochs were manually inspected and epochs with clear noise or movement artefacts were rejected.

#### Analysis of oscillatory power from the LFP

The LFP data was down sampled to 2,5 kHz during recording. The data was processed using custom made Python (v 3.11.4) programs, with the MNE (v. 1.4.2), Numpy (v. 1.25.2), Scipy (v. 1.14.1), Pandas (v.2.2.3) and and readSGLX from the SpikeGLX_Datafile_Tools package. The signals were low-pass filtered to 350 Hz and DC-components were removed by subtracting channel means. For each epoch, a time-voltage –plot covering all channels was plotted, and epochs with artifacts or noise were manually discarded based on the plotted figures.

Channels for analysis were selected on the basis of post-mortem histological investigation (i.e. recording sites of fluorescently marked electrodes). For the channels representing the LA, complex Morlet wavelet convolution was performed using the MNE tfr_array_morlet-function (mne.time_frequency.tfr_array_morlet), separately for theta (4 – 12 Hz), beta (13 – 30 Hz), low gamma (20 – 49 Hz) high gamma (50 – 90 Hz) and fast gamma (91 – 150 Hz) frequencies, and the mean power per animal on each frequency band was extracted for all epochs. The animal means at each frequency band were transferred to GraphPad Prism (v 10.3.1) for statistical analysis.

For power spectral densities, the power resulting from the Morlet wavelet convolution was obtained at 1 Hz steps from 4 Hz to 150 Hz separately, and normalized to the total power which was calculated using the composite trapezoidal rule (numpy.trapz). The mean and standard deviation of normalized power across animals in each group were calculated using numpy.mean ad numpy.std. The difference between movement and rest was calculated for each animal by calculating the mean absolute (non-normalized) power during rest and movement epochs, and subtracting the rest mean from the movement mean at each frequency separately.

#### Spike sorting and spike data analysis

The recordings were spike sorted with Kilosort 4 (Pachitariu et al., 2024) using default settings. Using open-source Python library phy (https://phy.readthedocs.io/en/latest/) (Rossant et al., 2016), good spike clusters with acceptable waveforms were first isolated, then their 2D principal component plots were inspected, and cross correlograms and the best channels were compared to split and merge the spike clusters as necessary.

Using custom made Python (v 3 11 4) programs, with the Scipy (v. 1.11.1; Virtanen et al., 2020) and scikit-learn (v 1 3 0; Pedregosa et al., 2011) packages, waveforms of the resulting spike clusters were parameterized five ways: (1) trough-to-peak time (Henze et al., 2000), (2) trough full width at half maximum (Weir et al., 2015), (3) 1^st^ and 2^nd^ peak-to-trough time asymmetry (Henze et al., 2002), (4) peak/trough amplitude ratio (Keshavarzi et al., 2022), (5) 1^st^ and 2^nd^ peak amplitude asymmetry (i.e. asymmetry index; Sirota et al., 2008). 20% of the spike clusters were manually classified as either wide or narrow waveform, and the remaining spike clusters were sorted using a linear discriminator trained on the 5 waveform parameters of the manually classified spike clusters.

Firing patterns of the spike clusters were parameterized seven ways: (1) number of spikes, (2) 95 percentile interspike interval, and five parameters derived from the auto-correlogram triple exponential curve fit (Petersen et al., 2021) (3) refractory period (4) asymptote (5) τ_burst_, (6) τ_decay_, (7) τ_rise_. Putative monosynaptic connections of the spike clusters were determined by convolving pairwise cross-correlograms with a partially hollowed gaussian kernel having a standard deviation of 10 ms and hollow fraction of 60% (Stark and Abeles, 2009) rather than employing a jitter resampling method. To determine a significant excitatory connection, a significant increase (p<0.001 expected from a Poisson process) in postsynaptic spike cluster firing lasting at least 0.4 ms must fall within a 0.4-4 ms window after presynaptic spike cluster firing (modified from English et al., 2017). Because inhibition tends to be less sharp than excitation, for a significant inhibitory connection, a significant decrease (p<0.05 expected from a Poisson process) in postsynaptic spike cluster firing lasting at least 0.99 ms falling within a 1-10 ms window after presynaptic spike cluster firing (modified from Headley 2021) was used as the criteria. Spike clusters originating both excitatory and inhibitory putative monosynaptic connections were classified ambiguous. Cross-correlograms of putative monosynaptic connections were all visually confirmed.

Combining plots of interspike interval histogram, auto-correlogram and putative monosynaptic connection, the 20% of the spike clusters with wide waveform were manually classified as either PN or wide-other spike clusters and similarly 20% of the spike clusters with narrow waveform were classified as either FSI or narrow-other spike clusters. The remaining spike clusters in both cases were sorted using a linear discriminator trained on the seven firing pattern parameters and putative monosynaptic connection of the manually classified spike clusters. All automatically classified spike clusters were all visually inspected and reclassified as necessary.

Validating our classification scheme, waveform linear discriminator routinely achieved discrimination score above 95%. While firing pattern linear discriminators were only able to achieve discrimination score above 75%, spike clusters originating inhibitory connections were always classified as FSIs or narrow-other.

Spike bursts of each PN and FSI spike cluster were determined by adopting the MaxInterval algorithm (Nex Technology, Colorado Spring, CO, USA), chosen based on a comparison of 8 burst detection algorithms (Cotterill et al., 2016). A burst starts when two consecutive spikes have an interspike interval less than 170 ms and ends when two consecutive spikes have an interspike interval more than 300 ms. A burst must contain at least three spikes and last at least 10 ms. Bursts with interburst interval less than 200 ms were merged into a single burst.

For spike-field coupling, the LFP phase during non-movement epochs was extracted for each channel and each central frequency using 1 Hz steps from 4 Hz to 150 Hz separately by Morlet wavelet convolution (mne.time_frequency.tfr_array_morlet). Pairwise phase consistency (PPC) values were calculated for each identified cluster as described in (Vinck et al., 2010) for each frequency, using the circular mean phase (scipy.stats.circmean) of the LFP from channels ± 7 channels away from the spike channel while excluding the 5 channels closest to the spike channel. Clusters with less than 10 spikes during the analyzed epochs were excluded from the PPC analysis. The frequencies yielding the highest PPC value in the theta range (4 – 12 Hz) and gamma range (20 – 150 Hz) were identified, and the Rayleigh test for circular non-uniformity (Wilkie., 1983) was used for the phase of each spike at those frequencies to determine whether the cluster showed significant phase locking. P-values less than 0.05 were considered significant. Spike probability distributions in 8 equally sized phase bins from 0 to 2π were calculated using the frequencies with highest PPC. Clusters showing uniform spike probability distribution across the LFP phase (p ≥ 0.05, Rayleigh test) were excluded from the distribution analysis.

